# Genetic and functional characterization of the natural transformation system in *Streptococcus constellatus*

**DOI:** 10.64898/2026.03.04.709537

**Authors:** Andreas Solberg Sagen, Kazi Shefaul Mulk Shawrob, Gabriela Salvadori, Roger Junges

**Affiliations:** Institute of Oral Biology, Faculty of Dentistry, University of Oslo, Oslo, Norway; Department of Public Health and Sport Sciences, Faculty of Social and Health Sciences, University of Inland Norway (INN), Elverum, Norway

**Author notes:** corresponding author, Roger Junges, DDS, PhD, Associate Professor, Institute of Oral Biology, Faculty of Dentistry, University of Oslo (UiO), Postboks 1052 Blindern 0316 - Oslo, Norway.

**Keywords:** DNA Transformation Competence, Streptococcus, Streptococcus constellatus, Gene Expression Regulation, Bacterial, Quorum Sensing, Regulon, Gene editing

## Abstract

*Streptococcus constellatus* is an opportunistic pathogen frequently associated with abscess formation in various body sites. While the species has been shown to acquire exogenous DNA through natural transformation, functional analyses of its underlying mechanisms and optimized genetic editing protocols remain limited. Thus, our aim was to characterize the natural transformation system in *S. constellatus* and investigate environmental factors regulating its natural transformation system. In addition, we sought to develop an optimized protocol for genome editing. Genomic analysis revealed that 73% of analyzed strains possess orthologs for essential competence regulon genes, with 55% harboring both a complete ComCDE-based operon and the necessary transformation machinery. While all complete genomes harbored three copies of the master regulator *sigX*, the accessory regulator *comW* was seemingly absent. Lacking the peptide exporter *comAB*, we demonstrated that *S. constellatus* utilizes the bacteriocin transporter *silED* for competence-stimulating peptide export. Gene expression assays indicated system activation at peptide concentrations as low as 4 nM, with peak *sigX* expression obtained over 60 nM. With the goal of optimizing gene editing strategies, we developed a protocol utilizing rich media supplemented with bovine serum albumin and calcium chloride, substantially increasing transformation frequencies. Furthermore, we observed that environmental stressors can upregulate the system, including hydrogen peroxide and subinhibitory concentrations of the antibiotics erythromycin, chloramphenicol, and ampicillin. Given the increasing clinical relevance of the anginosus group, elucidating horizontal gene transfer mechanisms can provide critical insights into the evolutionary success and pathogenic potential of these species.

## INTRODUCTION

*Streptococcus anginosus* group (SAG), comprises three different species of streptococci: *Streptococcus anginosus, Streptococcus intermedius*, and *Streptococcus constellatus*. Anginosus group species most notably colonize the oral cavity, throat, gastrointestinal and genital tracts as a commensal part of the human microbiome but can act as opportunistic pathogens causing non-invasive and invasive infections including pharyngitis, bacteraemia, and abscesses in various body sites [1]. The ability of anginosus group species to thrive in low-oxygen and acidic environments has been suggested to be an important adaptation supporting abscess formation, but they also present diverse defense mechanisms that can suppress the immune response [2,3]. Additionally, a large fraction of anginosus group genomes presents a variety of mobile genetic elements and prophages, likely contributing to rapid adaptability to external stimuli on a populational level [4–6]. In this context, the anginosus group, as a consequence of their plasticity and wide range of habitats in the human body, possesses a unique opportunity for horizontal gene transfer (HGT) across human microbiomes as they can interact with many species including priority human pathogens.

Anginosus species have been shown to naturally transform by incorporating exogenous DNA from the environment [7]. The natural transformation process and machinery relies on the ComDE two-component regulatory system in the mitis and anginosus group of streptococci, having most notably been studied in *Streptococcus pneumoniae* [8]. This system is based on a phosphorylase mechanism that is activated by the competence stimulating peptide (CSP), encoded by the gene *comC*, when binding extracellularly to the histidine kinase ComD. Upon the extracellular binding of the peptide, a phosphorylation cascade is initiated that activates the response regulator ComE [9]. Once activated, ComE triggers the expression of the alternative sigma factor SigX, which then enables the transcription of downstream genes involved in DNA uptake, recombination, DNA repair, stress-response, and fratricide [8]. Fratricide is a competence-associated mechanism which involves the lysis of neighbouring non-competent cells through the expression of specialized cell wall hydrolases [10]. These hydrolases are co-expressed with immunity factors that protect competent cells from autolysis, thereby ensuring selective targeting. The resulting release of extracellular DNA generates a local pool of genetic material that can be internalized and integrated via homologous recombination, functionally coupling cell lysis to horizontal gene transfer [11]. In *S. pneumoniae*, the competence system has been shown to affect gene expression of as much as 4% of the genome [12–14]. For the anginosus group, reports are more limited; however, previous studies in *S. anginosus* showed that strain SK52 contains most of the known essential competence genes and is capable of transforming in laboratory conditions [15]. In *S. constellatus*, an ortholog of *comCDE* and three copies of *sigX* have been identified previously [7,13]. In addition, a recent study corroborated the functional CSP sequence in the type-strain as a 16-amino acid peptide [16].

As interest in the anginosus group rapidly increases due to evidence of its virulence, such as abscess formation and potential tumor involvement [1,17], an expansion of the repertoire of genetic tools is likely to benefit the field. In addition, investigating HGT mechanisms in this group of species can provide novel insights into genome plasticity and survivability during the transition from commensal to pathogen. Here we show that the majority of *S. constellatus* genomes presents a complete competence regulon including the genes necessary for recombination. We further identified three copies of *sigX* in all completed genomes, with no *comW* present. Export of CSP relies on *silED* in this species, as the only transporter available for both CSP and SilCR, and we observed that *sigX* expression in response to synthetic CSP (sCSP) takes place already at low concentrations. Further, we present an optimized protocol for transformation of *S. constellatus* utilizing bovine serum albumin and calcium chloride that substantially improves transformation frequencies for both plasmid and amplicons used as donor DNA. Finally, through the assessment of a panel of environmental stressors, we observed that hydrogen peroxide (H_2_O_2_) and subinhibitory concentrations of the antibiotics erythromycin, chloramphenicol, and ampicillin stimulate *sigX* expression, which may have important implications for habitat adaptation in this species coupled with repercussions related to antibiotic selection for treatment of infections.

## MATERIALS AND METHODS

### Bacterial strains and culturing conditions

Bacterial strains, peptides, and antibiotics utilized in the study are listed in Table S1. Peptides and antibiotics were dissolved and stored according to the instructions of the manufacturer. *Streptococcus constellatus subsp. constellatus* type strain CCUG 24889 and isogenic strains were generally cultured in tryptic soy broth (TSB; Oxoid, Basingstoke, UK) at 37°C in a 5% CO_2_ humidified atmosphere. Brain-Heart Infusion (BHI; Oxoid, Basingstoke, UK) was prepared as described by the manufacturer, whereas THY was prepared from Todd-Hewitt Broth (THB; Oxoid, Basingstoke, UK) mixed with 0.2% yeast extract (Oxoid, Basingstoke, UK). Defined media CDM and C+Y_YB_ were prepared in accordance with the procedures described by Chang et al. (2011) and Stevens et al. (2011), respectively. Incubation of petri dishes was performed in anaerobic conditions for 48 hours as we observed more robust colonies, facilitating CFU determination. As described previously [20], bacterial pre-cultures for transformation and reporter assays were prepared by growing *S. constellatus* to optical density 600 (OD_600_) ≈0.4 from fresh colonies, supplemented with 15% sterile glycerol and stored at -20°C. When applicable, *S. anginosus* and *S. intermedius* strains were cultured using similar conditions.

### Reporter and mutant constructions

Mutants were constructed using overlapping PCR mutagenesis with sCSP as described previously by Junges et al. (2023). Oligonucleotides and donor DNA are listed in Table S2. Briefly, p*sigX* was amplified and fused with a firefly luciferase gene (*fluc*) and *aad9* (spectinomycin resistance cassette), originated from pFW5-luc [21], and inserted upstream of the p*sigX*_1_ locus in an amplicon. Subsequently, the amplicon was ligated via overlapping-PCR with two 2kb flanking regions p*sigX*_1_ to generate an amplicon for transformation into the chromosome of CCUG 24889, thus inserting the reporter locus upstream of the intact *sigX*_1_ locus. Deletion mutants were constructed similarly by amplifying and fusing 2-3 kbps flanking regions surrounding the deletion locus with overlapping-PCR. The final amplicon was then transformed into either CCUG 24889 or isogenic strains. Colonies for all mutants were then recovered in the appropriate selective antibiotic and further genotype confirmation was performed with PCR. DNA and plasmid extractions were performed with DNeasy blood and tissue kit (Qiagen, Hilden, Germany) and the ZymoPure Plasmid Miniprep Kit (Zymo Research, Irvine, CA, USA), respectively, at times with an additional lysis step to maximize yield. Fragment purification was conducted with MinElute PCR Purification Kit (Qiagen, Hilden, Germany).

### Transformation and reporter assays

Initial protocols were adapted from previous reports in other streptococci [15,22]. Transformation assays were performed by growing a 1:100 dilution of preculture to an early log-phase (OD_600_≈0.04), then donor DNA and 250 nM sCSP were added in 200 µL total volume. Cultures were generally incubated for 2 hours, then plated onto TSB agar with selective antibiotics and incubated for 48-72 hours in anaerobic conditions. Transformation frequency was calculated as the ratio of colony forming units (CFU) on selective agarose to CFU on non-selective agarose. Different conditions and parameters for transformation were assessed as described in the results section. As donor DNA, amplicon aSC011.3 and plasmid pRJ11, both conferring kanamycin resistance, were utilized generally. For reporter assays, pre-cultures of the reporters at OD_600_≈0.4 were diluted 1:100 in TSB with 100 nM D-luciferin (Synchem, Felsberg-Altenberg, Germany) in the presence or absence of different concentrations of sCSP at varying cell densities and antibiotic concentrations. Aliquots of 200 µL were distributed in 96-well microtiter plates. The cultures were monitored in a plate reader (Synergy HT; BioTek, Winooski, VT) for 24 hours at 37°C at high luminescence sensitivity. In post-run analysis the background values were subtracted, and luminescence (in RLU) was adjusted to the OD_600_ value recorded.

### Growth under stress conditions

Antibiotic stress assays were performed in 96-well microtiter plates incubated in a plate reader with 2-fold serial dilutions of spectinomycin, kanamycin, erythromycin, tetracycline, vancomycin, ciprofloxacin, chlorhexidine, rifampicin, chloramphenicol, novobiocin, or streptomycin. Initial concentrations are listed in Table S1. Similarly, the H_2_O_2_ experiment was performed using concentrations from 1 to 5 mM with dilutions freshly prepared from 3% concentrated H_2_O_2_. For testing the effects of pH, TSB solutions were adjusted prior to sterilization using HCl and NaOH, and recalibrated under sterile conditions after equilibration at 37°C. All experiments included a control without treatment compounds in addition to a blank medium control. Bacterial cultures were prepared from pre-cultures 1:100 and loaded onto each test group. Transformation assays were performed concomitantly in similar conditions using cultures with a OD_600_≈0.04 as described above. The results presented are representative of at least three independent biological experiments.

### RNA extraction, cDNA synthesis and RT-qPCR

Cells were harvested and RNA was extracted with a High-Pure RNA isolation kit (Roche) with an additional lysis step containing 10 mg/mL lysozyme, 100 U/mL mutanolysin, and 10mM Tris-HCL to maximize yield. Further, cDNA was synthesized using a First Strand cDNA synthesis kit (Thermo Scientific) following the manufacturer’s protocol with 1 µg RNA per reaction and an additional DNase treatment step. Gene expression was quantified using qPCR with Maxima SYBR Green/ROX (Thermo scientific) following manufacturer’s guidelines with 500 nM oligonucleotides, 10 ng cDNA and a three-step cycling protocol on an AriaMX Real-Time PCR system (Agilent, Santa Clara, USA). The primers were designed using NCBI Primer-BLAST with the relevant genes as template [23]. The quantification cycle (Cq) was normalized to the expression of a reference gene *gyrA* using Livak’s method [24].

### *In silico* regulon detection, bioinformatics and statistical analysis

Genomes of *Streptococcus constellatus* were obtained from the reference sequence database from NCBI on October 2024 and listed in Table S3 [25,26]. Gene orthologs of the competence regulon were identified using a combination of BLASTn and tBLASTn using gene queries listed in Table S4 [27]. The resulting hits were manually curated with particular focus to ensure inclusion of less conserved and smaller genes such as *silCR* and *comC* and remove duplicates from closely related genes such as *comE* and *silB* [28]. The 100 bp of the upstream region of genes in *S. constellatus* CCUG 24889 known to be regulated by SigX in *S. pneumoniae* were extracted and analyzed using Multiple Em for Motif Elicitation (MEME) v5.5, followed by fuzznuc to refine a more specific search pattern using previously known CIN-box motifs [14,29,30]. To search for additional binding sites of SigX in *S. constellatus* CCUG 24889 genome, EMBOSS fuzznuc was utilized with the refined search pattern. Guided by qPCR data, and an assembly of high-confidence motif binding sites, Find Individual Motif Occurrences (FIMO) was used to scan all the 33 genomes for high-confidence binding sites within 200 bp of a gene start codon [31]. The phylogenetic relationship between the strains of *S. constellatus* were determined using Phylogenomic Analyses for the Taxonomy and Systematics of Microbes (PHANTASM) with *S. pneumoniae* D39 as the outgroup [32]. Statistics and data visualization were either performed using R v4.4 or GraphPad Prism v10.2. For statistical analysis, the differences in treatment responses were analyzed using one-way repeated measures ANOVA or paired t-test across the changes in conditions. Significance level was set at 0.05.

## RESULTS

### The competence regulon in *S. constellatus*

The anginosus and the mitis group of streptococci present competence activation and bacteriocin production under regulation of the ComCDE and BlpRH operons, respectively [7,28]. In SAG, there is an analogous system of *blpA/B/C/H/R*, termed the *Streptococcus* Invasion Locus (Sil), coordinated by *silE/D/CR/B/A* [33]. The systems are structurally similar, and likely present a similar function [34]. Initially, we mapped the competence regulon in 33 genomes of *S. constellatus* deposited in the databases of the NCBI, utilizing the known regulon of *S. pneumoniae* and the reported genes from *S. anginosus* as reference [14,15]. Findings in *S. constellatus*, including a classification of the pheromones, identification of atypical features, and missing genetic elements clustered by genotypic similarity of the strains are summarized in Figure 1. Only one peptide cleavage/export ABC transporter system was identified across all the analyzed genomes. The locus was identified upstream of a gene cluster coding for bacteriocins and immunity proteins, a histidine kinase and a transcription factor, which resembles the BlpAB cluster from *S. pneumoniae*. Together with the closer sequence similarity with BlpAB from *S. pneumoniae* and SilDE from *S. anginosus* (Table S5), we putatively classify these genes as *silED* for *S. constellatus*.

**Fig 1.**
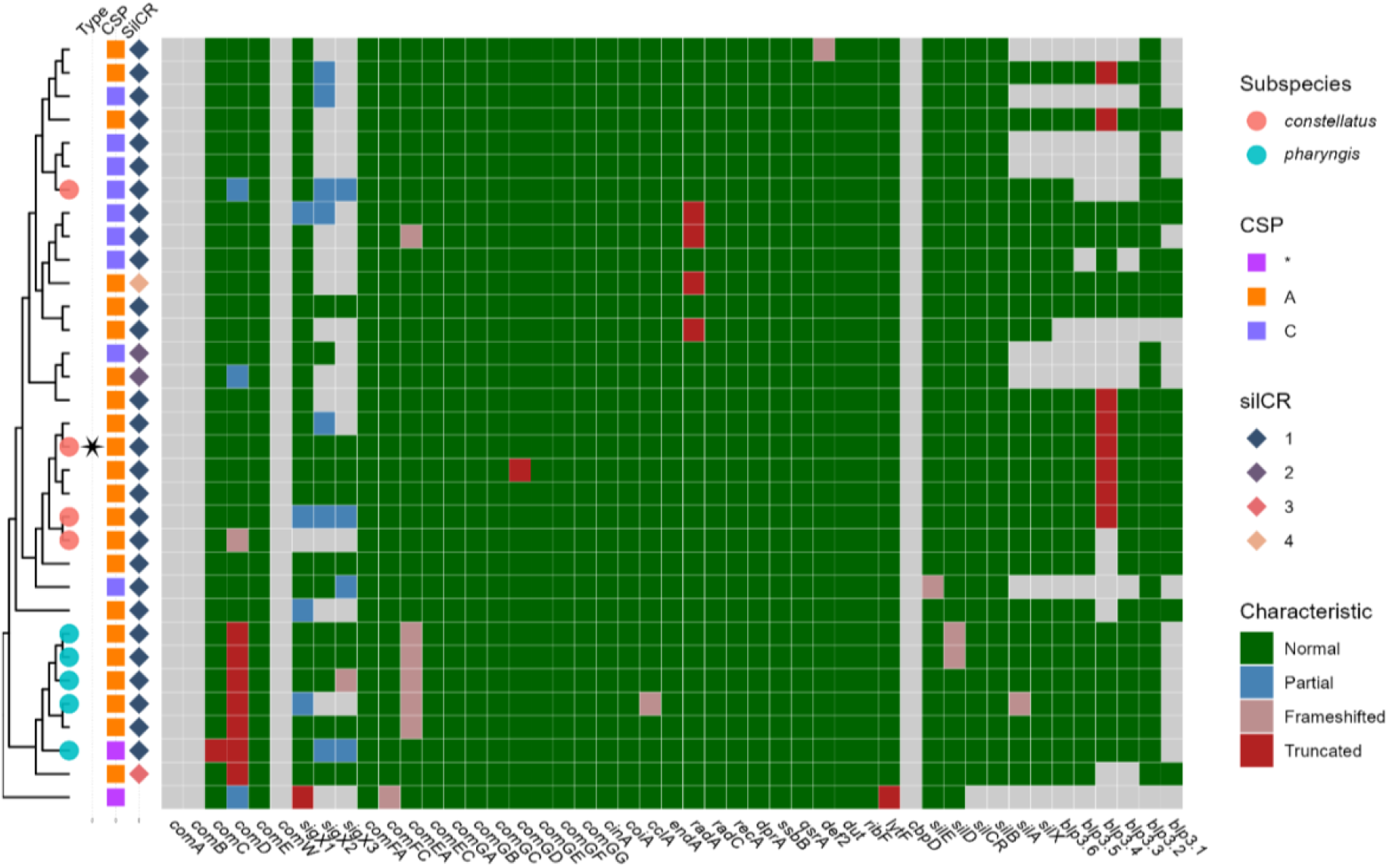
Distribution of competence-related genetic machinery in *S. constellatus*. The heatmap illustrates the status of genes involved in competence and fratricide systems. Gene characteristics are categorized as normal (green), partial (blue), frameshifted (brown), truncated (red), or absent (transparent). Phylogenetic analysis based on core genes of *S. constellatus* strains, annotated by subspecies and competence stimulating peptide (CSP) and SilCR isoforms. The type strain CCUG 24889 is indicated by a star.

Many strains of *S. constellatus subsp. pharyngis* presented a truncated *comD* and mutations leading to a frameshifted *comEA*. The truncation in *comD* takes place at the 3’-end of the gene, and a previous report shows that SigX can be activated despite the truncation. This is indicative that the lack of transformation observed for these strains is likely due to the frameshift in *comEA* [35]. It has previously been reported that species of the anginosus group have three copies of the *sigX* gene [13]. We found that 85% of the analyzed strains have at least one complete copy of *sigX* present. In the completely assembled genomes analyzed in this study, all carry three copies of the *sigX* gene, while the other strains that appear to lack three copies of *sigX* are draft assemblies. Thus, it is possible that the other copies are also present and functional but have not been reflected in the current sequencing data. If so, this is likely because of poor genome assembly due to the locus consisting of long repeating regions, particularly the 16S rRNA in the vicinity. In one of the genomes, *S. constellatus* subsp. *constellatus* SK53, no copy of *sigX* was identified, and one isolate *S. constellatus* SS_Bg39 presented only one partial copy of *sigX* at the edge of the contig. Further, the *sigX*-stabilizer *comW* was absent across the *S. constellatus* strains analyzed.

Two forms of putative mature CSP were identified based on *comC* sequences - DSRIRMGFDFSKLFGK and DRRDPRGIIGIGKKLFG - with the first sequence being more common and also found in the type-strain. *S. anginosus* does not present the murein hydrolase CbpD that is well-characterized in *S. pneumoniae*, but rather presents another peptidoglycan hydrolase named LytF, which has been suggested to be involved in fratricide in *S. gordonii* and *S. mutans* [11,36]. In all genomes surveyed of *S. constellatus*, an intact copy of *lytF* was identified, but not *cbpD* (Fig 1). In summary, we have identified that the competence system is conserved in at least 73% of the strains analysed and 55% of strains containing a complete regulon necessary for transformation.

### Identification of SigX binding motifs and putative regulon

After binding of sufficient levels of CSP, a phosphorylation cascade activates expression of *sigX*, which further activates late genes related to DNA incorporation, recombination, and fratricide [8]. Based on previous reports, a total of nine candidate genes were selected as part of the high confidence late competence regulon: *cclA, coiA, cinA, comEA, comFA, comGA, dprA, dut* and *ssbB* [14]. The 100 bp upstream regions of these genes were extracted and analyzed using MEME to identify conserved motifs. By doing this, we identified a conserved 10 bp motif, TTTRCGAATA. As expected, this sequence motif is similar to the 8 bp sequence, TACGAATA, consensus sequence found in *S. pneumoniae, S. mitis, S. pyogenes* and *S. infantarius* [14,37]. Additionally, we observed a thymine-rich upstream and adenine-rich downstream region in line with the extended CIN-box as reported by Slager et al. (2019). Utilizing the promoter sequences of the high-confidence SigX regulon described above, we refined an extended CIN-box motif for *S. constellatus* - WDBNNNHNNNHHYYNCGAATWDDND - and performed searches in the entire genome of the type strain. We combined results and identified several motifs upstream of genes or open reading frames within 100 bp, and some beyond, including *pheT, lytF, coaB, ciaH, murB*, and hypothetical protein (Fig 2A). Based on these searches, we assessed the expression of each gene with RT-qPCR with positive (250 nM sCSP) and negative (sterile water) controls in a *ΔcomC* background, incapable of activating *sigX* without external stimuli. Upon addition of sCSP, gene expression for most of the highlighted candidates was upregulated (Fig 2B). Of the genes with unclear involvement in natural transformation, *SCSC_RS09450* (referred to as hyp. prot in the figure) was highly upregulated, whereas both *pheT* and *ciaH* showed lower levels of upregulation. Slight upregulation was also observed in four binding sites located within gene boundaries, namely SCSC_RS05465 (1.30 ± 1.65), SCSC_RS05825 (2.63 ± 1.96), SCSC_RS05935 (2.41 ± 1.96) and SCSC_RS06910 (2.59 ± 2.61).

**Fig 2.**
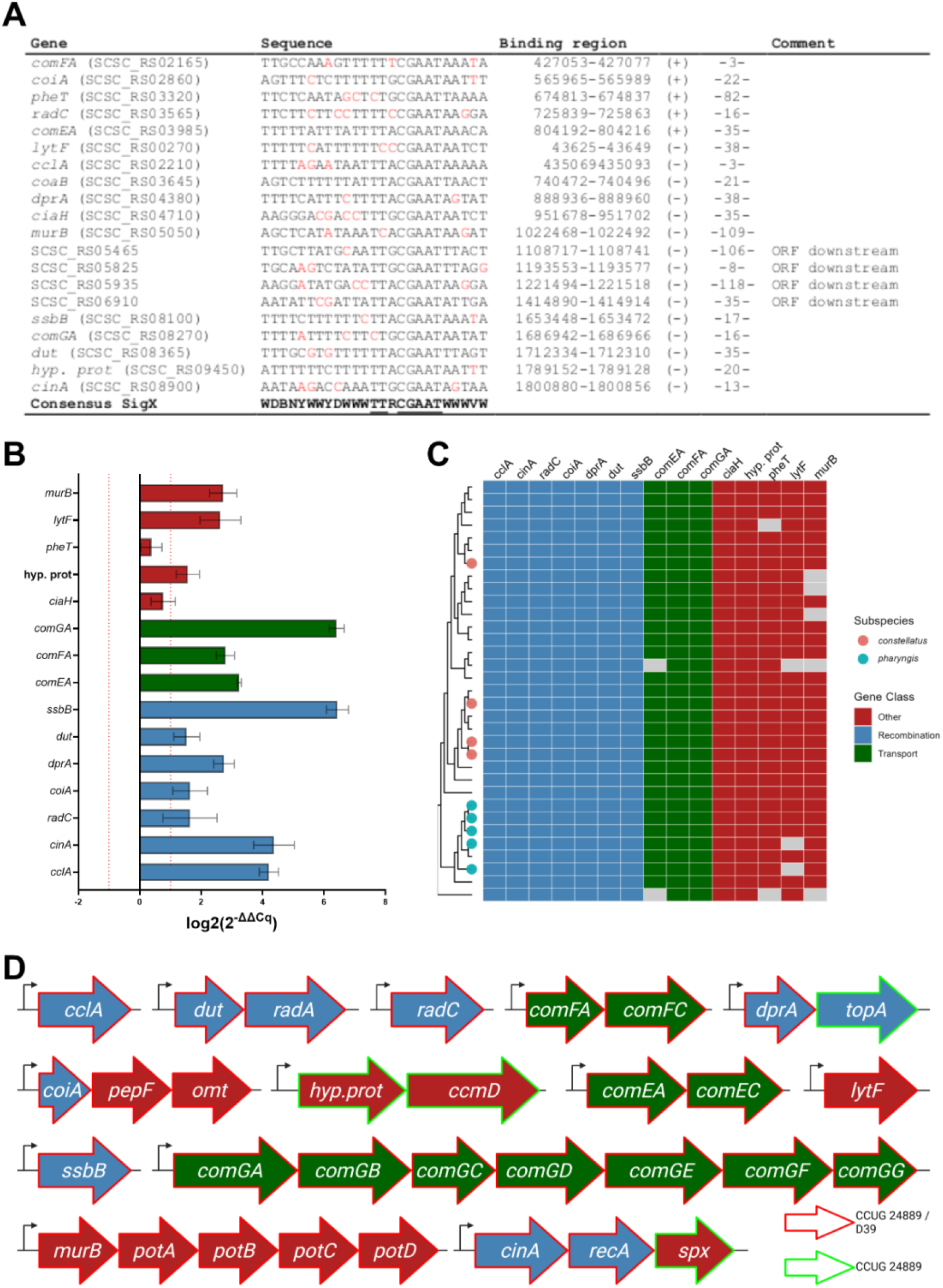
Characterization of the late competence regulon. **(A)** Alignment of the CIN-box motif and IUPAC consensus sequence upstream of predicted late genes. **(B)** RT-qPCR analysis of CSP-induced late gene expression. Data represent mean ± SEM for three biological replicates normalized to *gyrA*. **(C)** Analysis of CIN-box motifs identified across 33 genomes, categorized by functional class. **(D)** Genomic organization of gene clusters located downstream of identified CIN-boxes. Red outline indicates similar organization in type-strain CCUG 24889 and *S. pneumoniae* D39, whereas green outline indicates unique organization in *S. constellatus*.

We utilized these refined search patterns in FIMO to expand the search to 33 genomes of *S. constellatus* [29,31], and identified these genes to be mostly conserved similarly to the binding sites (Fig 2C). In terms of operon organization, *S. constellatus* appears to share the wider organization of *S. pneumoniae*. Comparing the *S. pneumoniae* D39 genome with *S. constellatus* CCUG 24889, we found the gene *spx* following *cinA* and *recA* in *S. constellatus* (Fig 2D), whereas it is located elsewhere in *S. pneumoniae*. The gene *spx* is a transcriptional regulator likely involved in stress responses [38]. Downstream of *dprA* in *S. constellatus* one can also identify a second gene, *topA*, which seemingly presents no homologous candidate in *S. pneumoniae*. Similarly, a CIN-box was identified upstream a unique hypothetical protein, with no homologous candidate in *S. pneumoniae*, followed by another gene *ccmD* also seemingly not present in *S. pneumoniae*.

### Kinetics of *sigX* expression in *S. constellatus*

While natural transformation has been traditionally studied in fast growing streptoccoci such as *S. pneumoniae* and *S. mutans, S. constellatus* exhibits growth dynamics characteristic of the broader anginosus group with a long lag-phase, followed by a slower doubling time [39]. This is observed in the laboratory in paralell to clinical conditions, where anginosys species are often observed to develop abscesses and chronic infections [40]. These characteristics require adaptation for the optimization of the existing protocols for competence and transformation studies. As a measurement of competence activity and with the goal of assessing the kinetics under different conditions, we constructed a luciferase reporter system in *S. constellatus* type strain as described previously by Junges et al. (2023). For the initial transformation assay of *S. constellatus* type strain CCUG 24889, we adapted from two protocols reported by Bauer et al. (2018) and Salvadori et al. (2017). Briefly, the type-strain was grown in TSB medium to OD_600_≈0.04, before 250 nM sCSP and 100 ng amplicon DNA were added. A number of colonies were obtained after 48 hours incubation in anaerobic conditions and a few putative mutants exhibiting the resistant phenotype were genotyped using colony PCR and functionally tested by assessment of luminescence emission (Fig 3A). We could observe a strong luminescence response in p*sigX-fluc* mutant in the presence of sCSP, but also without added peptide, potentially due to endogenous production of CSP. To confirm, a *comC* knockout mutant of the reporter was constructed. This Δ*comC* reporter did not show any activation compared to the background without addition of sCSP (Fig 3A), indicating that the reporter constructed in the type strain allowed for assessment of both endogenous production and response to exogenous stimuli.

**Fig 3.**
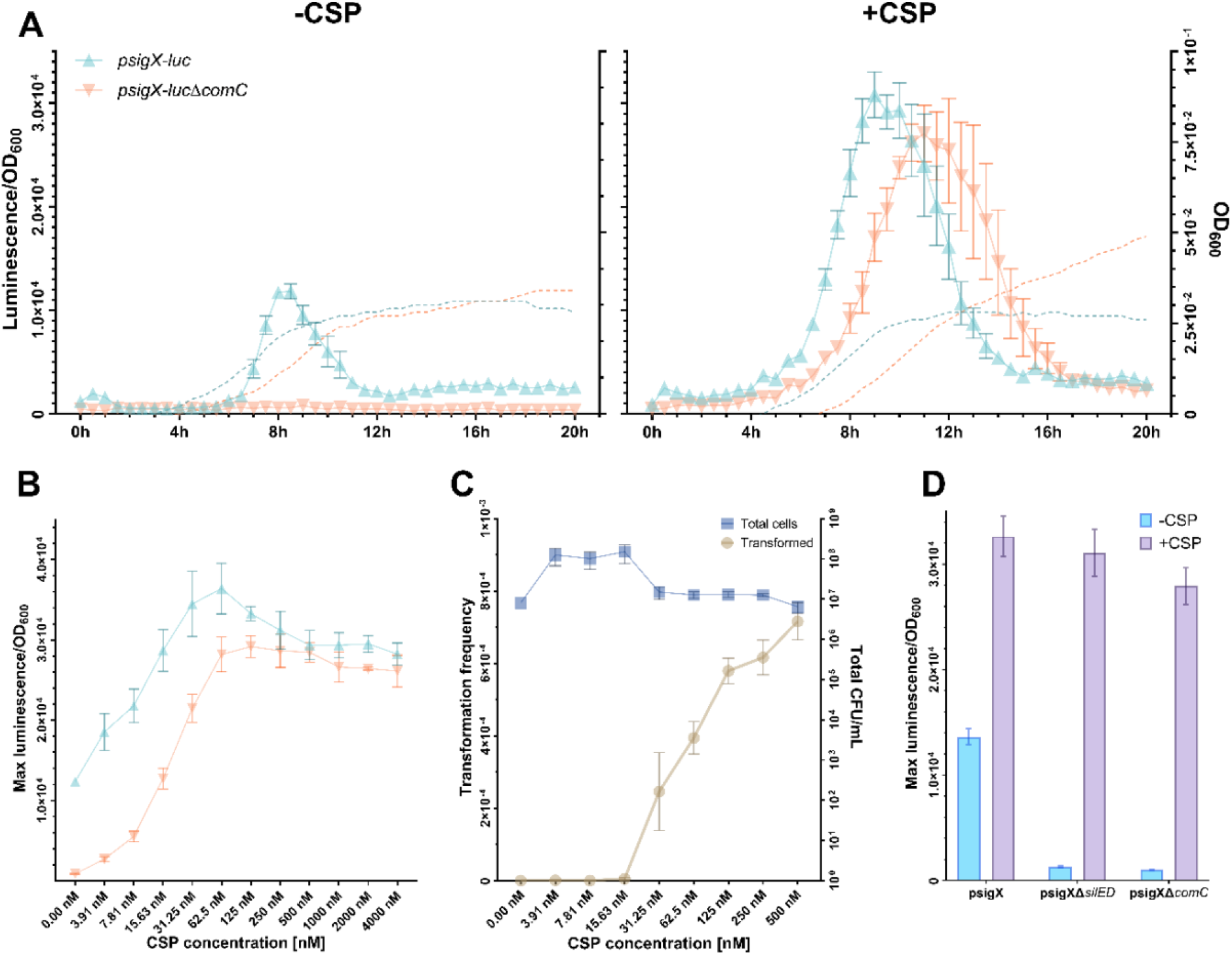
Parameters governing competence and natural transformation. **(A)** Temporal profiles of growth-adjusted luminescence (*sigX* expression) in the type strain and Δ*comC* mutant ± CSP. **(B)** Concentration curve of response to sCSP. **(C)** Dose-response of CSP concentration on maximum *sigX* expression and transformation frequency using a 3kb *kan*^*R*^ amplicon. **(D)** Impact of *silED* and *comC* deletions on endogenous and CSP-induced *sigX* expression. All data represent mean ± SEM of three independent biological experiments.

We further sought to reveal the response of *sigX* to different concentrations of sCSP (Fig 3B) and observed a statistically significant increase in *sigX* expression around 15 nM in the *comC* deficient mutant. While not statistically significant, an induced response was observed as low as 4 nM. The expression plateau was reached with 60 nM sCSP with no additional response on *sigX* expression observed for higher concentrations. Transformation frequency data indicate that a higher concentration of sCSP in the range between 125-500 nM yields higher transformation frequencies (Fig 3C). Finally, as we have identified in the first section of the results and as it has also been reported by others, only one peptide cleavage/export ABC transporter system is present in species of the anginosus group [33,34]. This system is comparatively more similar to BlpAB/SilED than to ComAB (Table S5). To understand if SilED is responsible for exporting CSP in this species and if there are any other redundant transporters, we constructed a *silED* knockout in the *S. constellatus* CCUG 24889 *psigX-fluc* strain. After knocking out *silED*, the endogenous expression of *sigX* was reduced to the same level of the Δ*comC* reporter (Figure 3D), indicating that SilED is responsible for exporting CSP with no other redundant exporters. In summary, we were able to assess both endogenous and exogenous assessment of CSP activity in *S. constellatus*. We further show that concentrations as little as 4 nm already activate the system in this species, and the minimum concentration required for maximum reporter activity is 60 nM. In addition, data indicate that *silED* is responsible for exporting CSP in *S. constellatus*.

### Optimizing conditions for competence and transformation

To gain insights into *S. constellatus* transformation and to optimize genetic editing protocols, we aimed to identify conditions that can affect competence development in this species. Optimal conditions for competence development can vary; however, growth medium and cell density have been previously reported as important components that can impact subsequent transformation frequencies significantly [22,41]. As such, assessment of *sigX* activity was performed using the reporter mutant grown in different media - from defined medium C+Y_YB_ and CDM, to complex medium THY, BHI and TSB. The addition of sCSP did not impact reporter activity in C+Y_YB_, CDM, or THY, but a difference was observed in TSB and BHI (Fig S1A). Transformation was tested in all the media listed and recovery of transformants was sucessful when utilizing TSB, in the conditions tested. It is possible that protocols for each medium would need to be optimized to obtain transformants; nevertheless, based on this finding and on the previous positive experiment outcomes for construction of reporters, we proceeded to use TSB as the method of choice given that it allowed transformation while also yielding a clear distinction in terms of *sigX* activation in the reporter with and without sCSP.

An important parameter in the optimization process was the initial dilution of the preculture. This factor is critical given that streptococci enter competence at a defined cell density [42]. To determine the optimal dilution, precultures frozen at early log phase were diluted 10-, 100-, or 1000-fold in TSB, with or without sCSP, and *sigX* expression was subsequently measured. Significant difference in reporter activity response was found depending on which dilution factor was chosen (Fig S1B). The endogenous response was seemingly stronger in cultures prepared with lower dilution factors, whereas the delta obtained from the addition of sCSP is more evident in higher dilutions. Based on these results and previous reports, a dilution factor of 1:100 was chosen to further assess the system [41].

Maximal *sigX* expression occurred between 6 to 10 hours after treatment with sCSP immediately after dilution. Based on our previous experience with transformation of *S. pneumoniae* [20,43], *S. mitis* [41], and *S. mutans* [20,44], we instead allowed the 1:100 diluted culture to grow to OD_600_≈0.04 prior to addition of sCSP and DNA, as this approach has previously resulted in significantly improved transformation frequencies. This growth stage typically takes 2-4 hours, however, once sCSP is added to a growing culture, peak *sigX* expression is reached within 2 hours (Fig 4A). To further develop the protocol, we asked whether the addition of calcium chloride (CaCl_2_) and bovine serum albumin (BSA), seen to improve transformation likely by assiting in DNA protection/stabilization and DNA-cell association, would affect transformation in *S. constellatus* [20,45,46]. We compared transformation frequencies in the presence of these compounds utilizing both amplicon and plasmid as donor DNAs. For amplicon DNA, transformation frequencies increased when adding 0.05% BSA (Fig 4B), but no discernible difference in combination with 0.5 mM CaCl_2_ was observed. For transformation with plasmid DNA, addition of a mix of 0.05% BSA and 0.5 mM CaCl_2_ increased the transformation frequency considerably, while any component on its own did not significantly change transformation yields. No significant increase in transformation frequency was observed when increasing concentrations to 5 mM CaCl_2_ or 0.5% BSA. As such, the optimal conditions for transformation were found to be with addition of 0.05% BSA and 0.5 mM CaCl_2_ when transforming with plasmid DNA and 0.05% BSA when utilizing amplicon DNA. In conclusion, we identified conditions in the protocol to transform *S. constellatus* that frequencies significantly. Further, we have successfully constructed mutants in all anginosus species utilizing this protocol (data not shown). Of note, while the length of the homogenous flanking loci can affect the transformation frequency [44], we did not observe any difference in transformation yields between a 3.5 kbp and a 7 kbp amplicon from the same genetic background. As such, most of the results presented are results of transformation with the former shorter amplicon.

**Fig 4.**
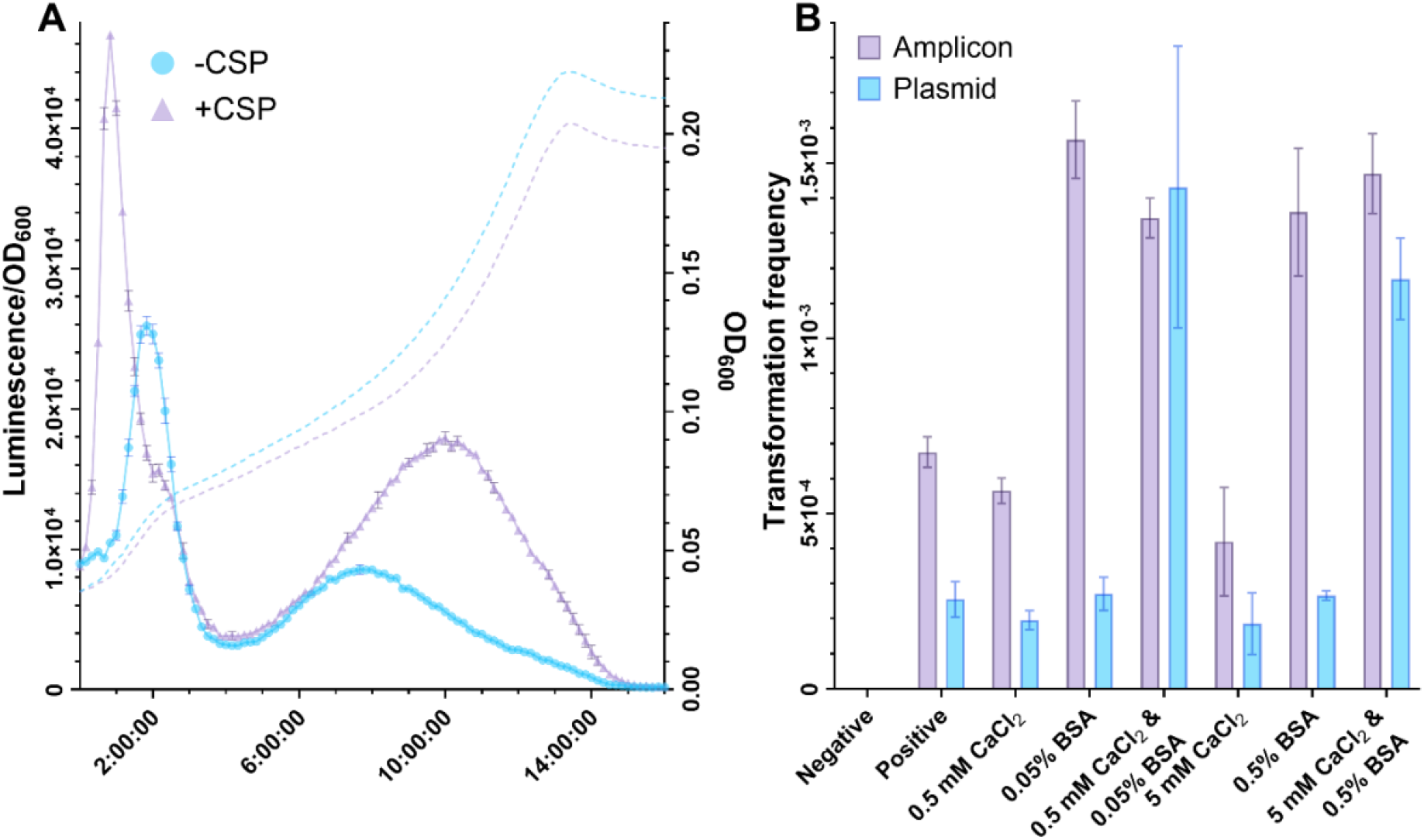
Optimization of the *S. constellatus* transformation protocol. **(A)** Impact of pre-growth inoculum density OD600 ≈ 0.04 on the timing of *sigX* expression. **(B)** Effect of BSA and CaCl_2_ on transformation frequencies for amplicon and plasmid DNA. All data represent mean ± SEM of three independent biological experiments.

### The effect of environmental stressors on natural transformation

Environmental stressors such as antibiotics have been shown to influence horizontal gene transfer and affect bacterial ability to enter the state of competence [47–49]. As the anginosus group of streptococci thrives at a variety of body habitats and challenging growth conditions such as tumour microenvironments and abscesses [50–52], we hypothesized that *S. constellatus* could present a distinct ability to respond to environmental stressors via the transformation system as an important vehicle for genetic diversity and DNA repair. We tested a range of antibiotics at different concentrations and identified that ampicillin, chloramphenicol and erythromycin generated an increased *sigX*-expression consistently at multiple subinhibitory concentrations (Fig 5). The strongest response on *sigX* expression was observed at 625 ng/mL for chloramphenicol and ampicillin, while erythromycin had its strongest response at 9.4 ng/mL. Most of the other antibiotics tested did not seemingly induce gene expression, and in some cases, e.g. novobiocin, seemed to show a repressive effect. For ampicillin and erytromycin that upregulated the system, we also observed higher transformation frequencies relative to the negative control, whereas transformation frequency did not appear to change with exposure to chloramphenicol. In conclusion, subinhibitory concentrations of ampicillin and erythromycin induced competence expression in *S. constellatus*, and led to phenotypically observed behavior. Further, as *S. constellatus* is often isolated from acidic and inflammatory environments, we tested transformation in a wide range of pH between 5.5 and 8.0 and were able to recover transformants between 6.5 and 8.0, with the highest frequencies between 7.0 and 7.5, as expected (Fig 6A). H_2_O_2_ is a reactive oxygen species and an important component in species-species competition of streptococci, as well as a potent antimicrobial component of the immune response. After exposure to subinhibitory concentrations of H_2_O_2_, we observed tolerance up to 3 mM H_2_O_2_, and an elevation of *sigX* expression at 0.5, 1, and 2 mM (Fig 6B). Additionally, higher transformation frequencies were also obtained in the presence of 1 and 2 mM H_2_O_2_ compared to the negative control. Altogether, these data indicate *S. constellatus* can adapt to diverse environmental conditions while maintaining activation of its competence system, which may provide competitive advantage under high stress environments.

**Fig 5.**
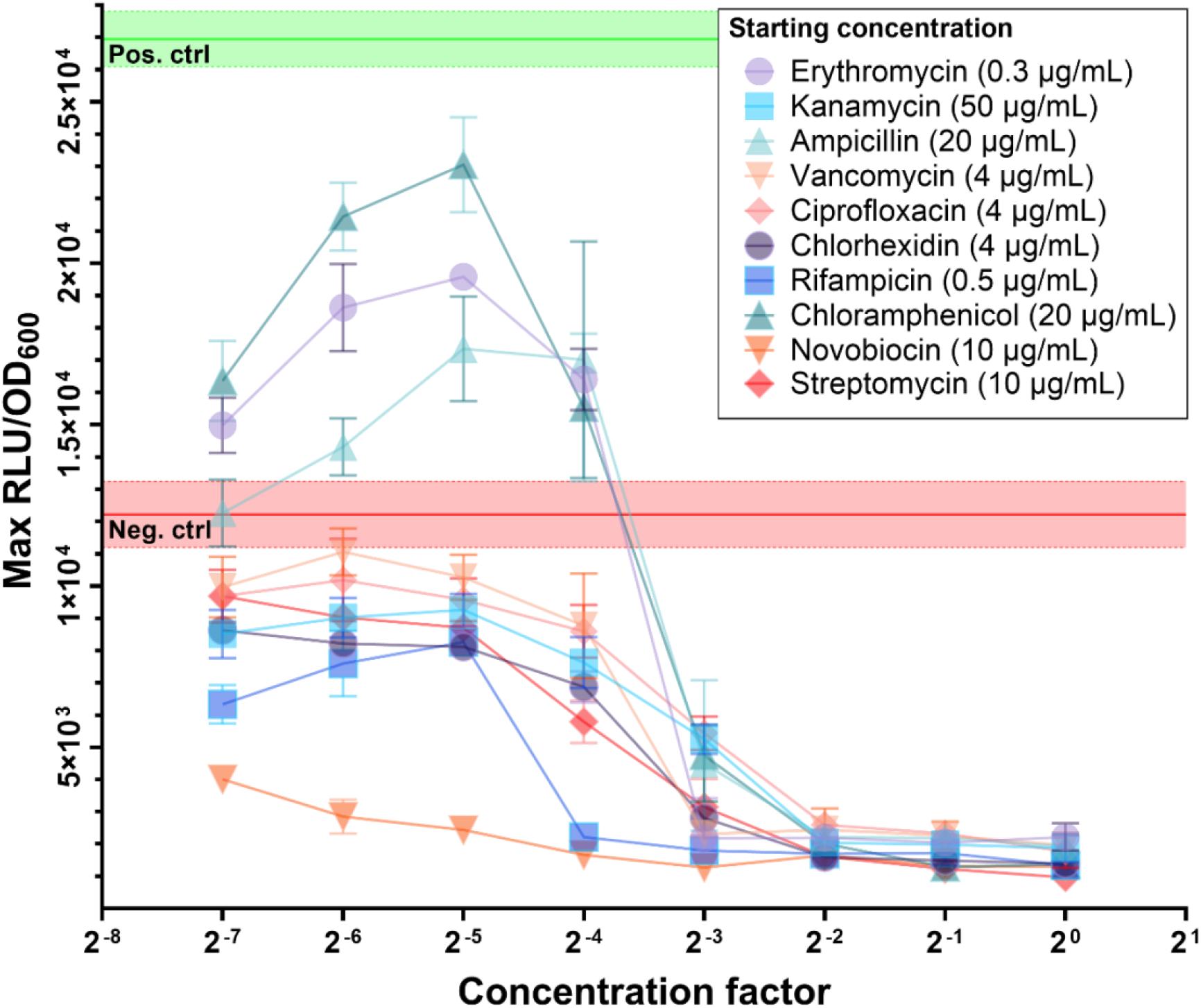
Response of *sigX* expression to antibiotic and antiseptic compounds. Maximum growth-adjusted luminescence of the *sigX* reporter strain (SC003) exposed to various antibiotics and antiseptics at inhibitory and sub-inhibitory concentrations. Green and red horizontal lines indicate negative (water) and positive (250 nM CSP) controls. All data represent mean ± SEM of three independent biological experiments.

**Fig 6.**
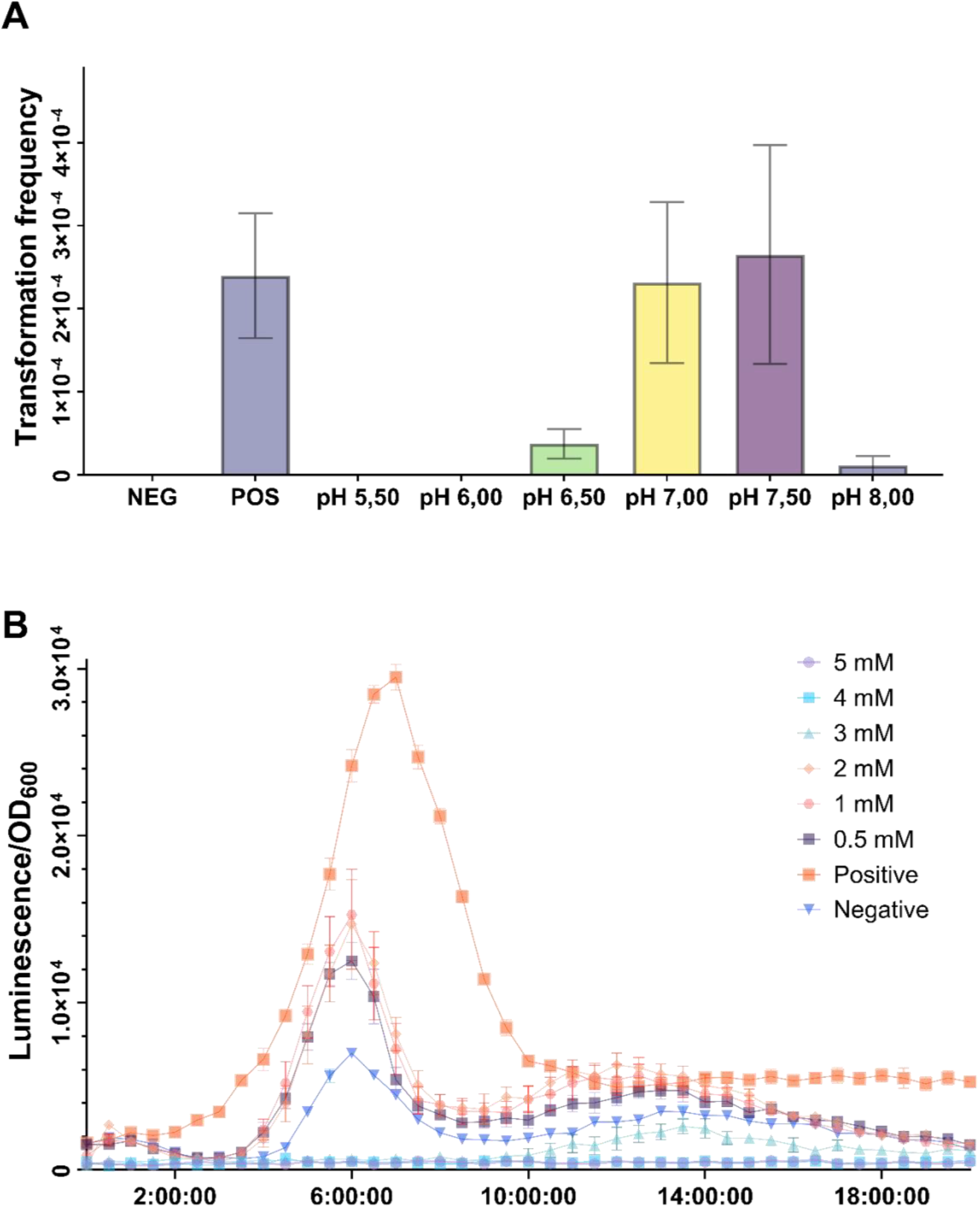
Environmental stressors effect on transformation and competence. **(A)** Effect of pH on transformation frequency in the type strain CCUG 24889. **(B)** Dose-dependent effect of growth-adjusted luminescence by hydrogen peroxide. All data represent mean ± SEM of three independent biological experiments. Negative (water) and positive (250 nM CSP) controls are represented.

## DISCUSSION

Horizontal gene transfer plays a central role in streptococcal adaptation, yet the mechanistic basis of natural transformation within the anginosus group remains limited. Although *S. constellatus* has previously been reported to undergo natural transformation, its regulatory circuit and genetic competence determinants have not been systematically characterized. In this study, we demonstrate that most *S. constellatus* strains harbor a complete competence regulon, including the core genes required for DNA uptake and homologous recombination. We have also shown that *S. constellatus*, and likely all species and subspecies in the anginosus group, rely on the bacteriocin exporter *silDE* for export of CSP. As adaptation to environmental challenges is clinically relevant for SAG, upregulation of the system and consequently an enhanced capability to transform was observed for *S. constellatus* in a variety of stress scenarios. Notably, subinhibitory antibiotic concentration of erythromycin, ampicillin, and chloramphenicol consistently activated *sigX* expression. Finally, with an optimized protocol, we increased transformation frequencies substantially, which further facilitates the construction of mutant strains and deepens our understanding of HGT in this important opportunistic pathogen.

While the interplay between competence and bacteriocin production is common among streptococci, the complete absence of ComAB and consequently reliance on SilDE to activate competence seems to be a unique feature of the anginosus group. While ComAB is generally conserved across all pneumococcal strains and essential for competence regulation, about a fourth of the strains also encode an intanct BlpAB, analogous to SilDE [33]. Interestingly, it has been shown that in BlpAB-positive strains of pneumococcus, the BlpAB transporter can mediate the secretion of both bacteriocins and CSP, suggesting that these strains could activate competence independently of ComAB [33]. In addition, Kjos et al. (2016) have demonstrated that when BlpAB is disrupted, ComAB can compensate by exporting BlpC. These findings support the notion of redundant function between the transporters ComAB and BlpAB, as observed in our results indicating that *silED* is essential for exporting CSP in *S. constellatus*. In the pneumococcus, it was further observed that strains containing both active transporters seem to have a competitive advantage, likely acting as agressors to outcompete other strains and species [33]. Other streptococcal species such as *Streptococcus thermophilus* and *Streptococcus gallolyticus* have been shown to contain functional BlpRH systems that regulate bacteriocin production through quorum sensing [28,54]. The presence of a single transporter in SAG can also have implications into its survival against other species in human microbiomes, and further research into competition assays in clinically-relevant scenarios is warranted.

No ComW orthologs were identified in *S. constellatus* in this study. In *S. pneumoniae*, ComW seemingly presents two main functions: stabilizing SigX and enhancing its ability to form an active RNA polymerase holoenzyme complex, possibly by assisting the alternative sigma factor in the competition against with the primary sigma factor for RNA polymerase binding [55]. It has also been shown that ComW binds to DNA, though not in a sequence-specific way, and this DNA-binding activity appears essential for its function in transformation [56]. While *comW* seems to be conserved across species of the mitis group, including *S. pneumoniae, S. mitis*, and *S. oralis*, previous reports have not identified orthologs in *S. anginosus* [15]. This goes in line with the findings we presented here regarding *S. constellatus* genomes. We did, however, noticed one study reporting a possible *comW* in *S. anginosus* SK1138 [56]. We expanded the search to other anginosus species, namely *S. intermedius* and *S. anginosus*, utilizing the SK1138 gene and identified a total of five strains of *S. anginosus* with an ortholog *comW*. No matches had been obtained with the *comW* gene from *S. pneumoniae* D39. Pairwise comparison of the ComW proteins in *S. pneumoniae* D39 and *S. anginosus* SK1138 showed 40% identity and 57.5% similarity. As a follow-up, homology searches across the domain of bacteria, excluding SAG species, showed a close similarity between the *comW* variants found in the SAG species with variants found in four different strains of *Streptococcus cristatus*. While the study of mechanistics of *comW* in this strain was outside the scope of this study, we were able to recover transformants in one of the *comW*(+) strains with similar yields compared to *comW*(-) strains (data not shown). Finally, another study identified mutations in the primary sigma factor, encoded by *rpoD*, that would bypass the need for *comW* [57]. However, such mutations were not present in the *S. constellatus* genomes assessed in this study. For the SigX-regulon, based on the identification of bindings sites, while the organization in *S. constellatus* is generally similar to *S. pneumoniae*, there were a variety of genes or ORFs with no orthologs identified and with unknown functions in the competence development process. These could be targets for further investigation and the strategy we employed for the motif-pattern search may be applicable in other bacteria, as most of the targets identified showed upregulation in the system upon sCSP treatment. Finally, the coupling with transcriptome data may be a useful resource in such cases.

A recent study confirmed the sequence of the *S. constellatus* CSP as a 16-amino acid peptide (DSRIRMGFDFSKLFGK) and explored the residue significance via the construction of peptide analogs [16]. With structure-activity assays using alanine and D-amino acid scans, the authors revealed that the N-terminal region is vital for activation of the system, and, notably, the C-terminal region is also crucial for *S. constellatus*. As such, they were able to identify the analog CSP I4A to be a more potent activator of the competence system. However, this increased response and potency was not translated into improved transformation frequencies. Further, the successful identification of CSP in supernatants indicates the functionality of the system and goes in line with our findings suggesting that the natural transformation system of *S. constellatus* plays a role *in vivo*. Also similar to our findings was their discovery that low concentrations of CSP were observed to upregulate the system in their reporter system. The authors observed that the competence regulon had no role in biofilm formation, in contrast to previous observations in other streptococci [58,59]. Increased biofilm formation is likely due to cell lysis and release of eDNA. On this note, we identified that the murein hydrolase LytF was present in all genomes surveyed of *S. constellatus* as part of the SigX-regulon. Despite copies of LytF being identified in a variety of streptococcal species, e.g. *S. gordonii, S. sanguinis, S. infantarius* and *S. cristatus*, their structure and loci can diverge across species [11]. While *S. gordonii* DL1 has been shown to possess a LytF with three BSP domains followed by a CHAP domain, *S. sanguinis* SK36 presents a larger gene with five BSP domains [60]. The structure and loci of the LytF in *S. constellatus* is similar to *S. sanguinis* SK36. Interestingly, a recent study in SK36 reported findings supporting the role of LytF in extrusion of the transformation pilus across the cell wall, and not directly on fratricide [61]. The authors also investigated the role of a CAAX protease as a potential LytF-immunity gene but found no evidence of increased lysis in knockout mutants. In *S. constellatus*, this protease is located in the Sil locus and named *silX*. Its function has been suggested to involve the export and processing of the peptide SilCR, while no immunity has been detected towards angicin [62]. In *S. mutans*, it has been recently reported that LytF plays a role in the release of cytoplasmatic membrane vesicles via cell wall degradation [61], also resulting in lysis of a subpopulation of cells [63]. One interesting observation in our study was the absence of the typical growth arrest following sCSP addition, which is generally attributed to lysis of a subpopulation of cells during competence [10], and under the conditions tested, this was not observed in *S. constellatus*. This lack of detectable lytic activity may support the previous findings in *S. sanguinis*, suggesting that the lytic potential of *lytF* could be condition dependent.

Environmental stressors have been shown to influence competence in a variety of bacterial species including *S. pneumoniae*, and *Legionella pneumophila* [49,64]. In *S. constellatus*, we identified that subinhibitory concentrations of erythromycin, chloramphenicol, and ampicillin induced *sigX* expression, with increased transformation rates observed with erythomycin and ampicillin. Antibiotics that target protein synthesis, such as erythromycin and chloramphenicol, can lead to a higher number of misfolded proteins, which in turn can reduce CSP degradation extracellularly given the capacity of the HtrA protease for target processing [19]. As such, the stronger activation of *sigX* in these two cases may be due to CSP protection extracelluarly; however, it is interesting that higher transformation rates were observed for erythromycin but not chloramphenicol. The lack of effect on transformation rates despite increased *sigX* expression has been observed previously in the case of erythromycin and tetracycline [65], for instance, and may be a result of a general stress response affecting gene transcription or interference with downstream protein synthesis in the conditions that were tested. While ampicillin was not seen to stimulate competence previously in *S. pneumoniae* [49,65], we observed an inducing effect in *S. contellatus*. As beta-lactams interfere with the cell wall, it is a possibility that the presence of ampicillin at subinhibitory concentrations can lead to pore formation facilitating the translocation of proteins/peptides between intra-and extraceullar environments. As antibiotics that affect DNA replication elongation, e.g. novobiocin, ciprofloxacin, have been shown to induce competence via an increase of the *comCDE* gene dosage that are located close to the replication origin in *S. pneumoniae*, we also expected to see induction of *sigX* by these compounds [66]. However, no stimulation was observed with a seemingly repression of *sigX* when cells were exposed to novobiocin. In addition, while kanamycin has been shown to be a potent inducer of competence in *S. pneumoniae*, no *sigX* upregulation was observed in our study [66]. As antibiotics acting through similar pathways can affect streptococci divergently, this supports the notion that their effect is potentially species-dependent and further research is warranted to better unravel the dynamics of HGT in a community level under stress.

Altogether, the findings of this study expand our knowledge on natural transformation in streptococci and further faciltiate the study of molecular mechanisms of pathogenicity in the anginosus group through the development of a genome editing protocol with higher transformation frequencies. In addition, data on environmental stressors influencing the rate of HGT is particularly relevant for the study of adaptability and survival mechansims in this group of species. These findings may also carry implications clinically as (i) these species can be exposed to a variety of antibiotic formulations and concentrations as part of the commensal microbiota, and (ii) as some of these antibiotics can be recommended for the treatment of streptococcal infections. For instance, beta-lactams and ampicillin are part of the line of treatment as viable options against infective endocarditis caused by viridans group and other types of streptococcal infections [67–70]. Thus, understanding the mode through which different antimicrobial drugs and formulations within the same class and mode of action divergently affect bacterial physiology and resistance rates is a priority.

## ACKNOWLEDGEMENTS

We are grateful to Prof. Don Morrison (UIC Biological Sciences, Chicago, IL, USA) for valuable discussions on experimental ideas and data interpretation. We would also like to thank Heidi Aarø Åmdal and Ali-Oddin Naemi for technical assistance in the laboratory.

## AUTHOR CONTRIBUTIONS

A.S.S., K.S.M.S., G.S., R.J.: study concept and design; A.S.S., K.S.M.S.: acquisition of data; A.S.S., K.S.M.S., G.S., R.J.: analysis and interpretation of data; A.S.S., K.S.M.S., R.J.: drafting of the manuscript; A.S.S., K.S.M.S., G.S., R.J.: critical revision of the manuscript; A.S.S., K.S.M.S.: statistical analysis; R.J.: study supervision. All authors read and approved the final manuscript.

## REFERENCES

1. Asam D, Spellerberg B. Molecular pathogenicity of Streptococcus anginosus. Molecular Oral Microbiology. 2014;29(4):145–55. doi:10.1111/omi.12056

2. Sasaki M, Kodama Y, Shimoyama Y, Ishikawa T, Kimura S. Aciduricity and acid tolerance mechanisms of Streptococcus anginosus. The Journal of General and Applied Microbiology. 2018;64(4):174–9. doi:10.2323/jgam.2017.11.005

3. Sitkiewicz I. How to become a killer, or is it all accidental? Virulence strategies in oral streptococci. Molecular Oral Microbiology. 2018;33(1):1–12. doi:10.1111/omi.12192

4. Nguyen SV, McShan WM. Chromosomal islands of Streptococcus pyogenes and related streptococci: molecular switches for survival and virulence. Front Cell Infect Microbiol. 2014;4. doi:10.3389/fcimb.2014.00109

5. Shawrob KSM, Sagen AS, Lunde TM, Sribasgaran B, Ghafoor S, Al-Haroni M, et al. Mapping antibiotic resistance determinants in oral streptococci. BioRxiv. 2026. doi:10.64898/2026.02.06.700101

6. Wang Y, Liu T, Sida Y, Zhu Y. Diversity and Evolution of the Mobilome Associated with Antibiotic Resistance Genes in Streptococcus anginosus. Microbial Drug Resistance. 2025;31(2):52–63. doi:10.1089/mdr.2024.0229

7. Håvarstein LS, Hakenbeck R, Gaustad P. Natural competence in the genus Streptococcus: evidence that streptococci can change pherotype by interspecies recombinational exchanges. Journal of Bacteriology. 1997;179(21):6589–94. doi:10.1128/jb.179.21.6589-6594.1997

8. Salvadori G, Junges R, Morrison DA, Petersen FC. Competence in Streptococcus pneumoniae and Close Commensal Relatives: Mechanisms and Implications. Front Cell Infect Microbiol. 2019;9. doi:10.3389/fcimb.2019.00094

9. Pestova EV, Håvarstein LS, Morrison DA. Regulation of competence for genetic transformation in Streptococcus pneumoniae by an auto-induced peptide pheromone and a two-component regulatory system. Molecular Microbiology. 1996;21(4):853–62. doi:10.1046/j.1365-2958.1996.501417.x

10. Johnsborg O, Eldholm V, Bjørnstad ML, Håvarstein LS. A predatory mechanism dramatically increases the efficiency of lateral gene transfer in Streptococcus pneumoniae and related commensal species. Molecular Microbiology. 2008;69(1):245–53. doi:10.1111/j.1365-2958.2008.06288.x

11. Berg KH, Ohnstad HS, Håvarstein LS. LytF, a Novel Competence-Regulated Murein Hydrolase in the Genus Streptococcus. Journal of Bacteriology. 2012;194(3):627–35. doi:10.1128/jb.06273-11

12. Lee MS, Morrison DA. Identification of a New Regulator in Streptococcus pneumoniae Linking Quorum Sensing to Competence for Genetic Transformation. Journal of Bacteriology. 1999;181(16):5004–16. doi:10.1128/jb.181.16.5004-5016.1999

13. Olson AB, Kent H, Sibley CD, Grinwis ME, Mabon P, Ouellette C, et al. Phylogenetic relationship and virulence inference of Streptococcus Anginosus Group: curated annotation and whole-genome comparative analysis support distinct species designation. BMC Genomics. 2013;14(1):895. doi:10.1186/1471-2164-14-895

14. Slager J, Aprianto R, Veening JW. Refining the Pneumococcal Competence Regulon by RNA Sequencing. Journal of Bacteriology. 2019;201(13):10.1128/jb.00780-18. doi:10.1128/jb.00780-18

15. Bauer R, Mauerer S, Grempels A, Spellerberg B. The competence system of Streptococcus anginosus and its use for genetic engineering. Molecular Oral Microbiology. 2018;33(2):194–202. doi:10.1111/omi.12213

16. Mehrani M, Mull RW, Guo M, Renshaw CP, Singh RJ, Tal-Gan Y. Deciphering the Regulatory Role and Molecular Interactions That Drive the Competence Regulon Quorum Sensing Circuitry in Streptococcus constellatus. Biochemistry. 2025. doi:10.1021/acs.biochem.5c00195

17. Fu K, Cheung AHK, Wong CC, Liu W, Zhou Y, Wang F, et al. Streptococcus anginosus promotes gastric inflammation, atrophy, and tumorigenesis in mice. Cell. 2024;187(4):882-896.e17. doi:10.1016/j.cell.2024.01.004 PubMed PMID: 38295787.

18. Chang JC, LaSarre B, Jimenez JC, Aggarwal C, Federle MJ. Two Group A Streptococcal Peptide Pheromones Act through Opposing Rgg Regulators to Control Biofilm Development. PLOS Pathogens. 2011;7(8):e1002190. doi:10.1371/journal.ppat.1002190

19. Stevens KE, Chang D, Zwack EE, Sebert ME. Competence in Streptococcus pneumoniae Is Regulated by the Rate of Ribosomal Decoding Errors. mBio. 2011;2(5):10.1128/mbio.00071-11. doi:10.1128/mbio.00071-11

20. Junges R, Khan R, Tovpeko Y, Åmdal HA, Petersen FC, Morrison DA. Markerless Genome Editing in Competent Streptococci. In: Seymour GJ, Cullinan MP, Heng NCK, Cooper PR, editors. Oral Biology: Molecular Techniques and Applications [Internet]. New York, NY: Springer US; 2023 [cited 2024]. p. 201–16. Available from: https://doi.org/10.1007/978-1-0716-2780-8_13 doi:10.1007/978-1-0716-2780-8_13

21. Podbielski A, Woischnik M, Leonard BAB, Schmidt KH. Characterization of nra, a global negative regulator gene in group A streptococci. Molecular Microbiology. 1999;31(4):1051–64. doi:10.1046/j.1365-2958.1999.01241.x

22. Salvadori G, Junges R, Khan R, Åmdal HA, Morrison DA, Petersen FC. Natural Transformation of Oral Streptococci by Use of Synthetic Pheromones. In: Seymour GJ, Cullinan MP, Heng NCK, editors. Oral Biology: Molecular Techniques and Applications [Internet]. New York, NY: Springer; 2017 [cited 2025]. p. 219–32. Available from: https://doi.org/10.1007/978-1-4939-6685-1_13 doi:10.1007/978-1-4939-6685-1_13

23. Ye J, Coulouris G, Zaretskaya I, Cutcutache I, Rozen S, Madden TL. Primer-BLAST: A tool to design target-specific primers for polymerase chain reaction. BMC Bioinformatics. 2012;13(1):134. doi:10.1186/1471-2105-13-134

24. Livak KJ, Schmittgen TD. Analysis of Relative Gene Expression Data Using Real-Time Quantitative PCR and the 2-ΔΔCT Method. Methods. 2001;25(4):402–8. doi:10.1006/meth.2001.1262

25. O’Leary NA, Cox E, Holmes JB, Anderson WR, Falk R, Hem V, et al. Exploring and retrieving sequence and metadata for species across the tree of life with NCBI Datasets. Sci Data. 2024;11(1):732. doi:10.1038/s41597-024-03571-y

26. Pruitt KD, Tatusova T, Maglott DR. NCBI Reference Sequence (RefSeq): a curated non-redundant sequence database of genomes, transcripts and proteins. Nucleic Acids Res. 2005;33(Suppl_1):D501–4. doi:10.1093/nar/gki025

27. Altschul SF, Gish W, Miller W, Myers EW, Lipman DJ. Basic local alignment search tool. Journal of Molecular Biology. 1990;215(3):403–10. doi:10.1016/S0022-2836(05)80360-2

28. Dhaked HPS, Biswas I. Distribution of two-component signal transduction systems BlpRH and ComDE across streptococcal species. Front Microbiol. 2022;13. doi:10.3389/fmicb.2022.960994

29. Bailey TL, Williams N, Misleh C, Li WW. MEME: discovering and analyzing DNA and protein sequence motifs. Nucleic Acids Research. 2006;34(Suppl_2):W369–73. doi:10.1093/nar/gkl198

30. Rice P, Longden I, Bleasby A. EMBOSS: The European Molecular Biology Open Software Suite. Trends in Genetics. 2000;16(6):276–7. doi:10.1016/S0168-9525(00)02024-2

31. Grant CE, Bailey TL, Noble WS. FIMO: scanning for occurrences of a given motif. Bioinformatics. 2011;27(7):1017–8. doi:10.1093/bioinformatics/btr064

32. Wirth JS, Bush EC. Automating microbial taxonomy workflows with PHANTASM: PHylogenomic ANalyses for the TAxonomy and Systematics of Microbes. Nucleic Acids Research. 2023;51(7):3067–77. doi:10.1093/nar/gkad196

33. Wang CY, Patel N, Wholey WY, Dawid S. ABC transporter content diversity in Streptococcus pneumoniae impacts competence regulation and bacteriocin production. Proceedings of the National Academy of Sciences. 2018;115(25):E5776– 85. doi:10.1073/pnas.1804668115

34. Mendonca ML, Szamosi JC, Lacroix AM, Fontes ME, Bowdish DM, Surette MG. The sil Locus in Streptococcus Anginosus Group: Interspecies Competition and a Hotspot of Genetic Diversity. Front Microbiol. 2017;7. doi:10.3389/fmicb.2016.02156

35. Lacroix AMG. Investigation of Competence Heterogeneity in Streptococcus Milleri Group Clinical Isolates [thesis] [Internet]. 2014 [cited 2024]. Available from: https://macsphere.mcmaster.ca/handle/11375/14246

36. Xu Y, Kreth J. Role of LytF and AtlS in eDNA Release by Streptococcus gordonii. PLOS ONE. 2013;8(4):e62339. doi:10.1371/journal.pone.0062339

37. Morrison DA, Guédon E, Renault P. Competence for Natural Genetic Transformation in the Streptococcus bovis Group Streptococci S. infantarius and S. macedonicus. Journal of Bacteriology. 2013;195(11):2612–20. doi:10.1128/jb.00230-13

38. Turlan C, Prudhomme M, Fichant G, Martin B, Gutierrez C. SpxA1, a novel transcriptional regulator involved in X-state (competence) development in Streptococcus pneumoniae. Molecular Microbiology. 2009;73(3):492–506. doi:10.1111/j.1365-2958.2009.06789.x

39. Fei F, Mendonca ML, McCarry BE, Bowdish DME, Surette MG. Metabolic and transcriptomic profiling of Streptococcus intermedius during aerobic and anaerobic growth. Metabolomics. 2016;12(3):46. doi:10.1007/s11306-016-0966-0

40. Al Majid F, Aldrees A, Barry M, Binkhamis K, Allam A, Almohaya A. Streptococcus anginosus group infections: Management and outcome at a tertiary care hospital. Journal of Infection and Public Health. 2020;13(11):1749–54. doi:10.1016/j.jiph.2020.07.017

41. Salvadori G, Junges R, Morrison DA, Petersen FC. Overcoming the Barrier of Low Efficiency during Genetic Transformation of Streptococcus mitis. Front Microbiol. 2016;7. doi:10.3389/fmicb.2016.01009

42. Gaustad P, Morrison DA. Induction of transformation in streptococci by synthetic competence stimulating peptides. Methods Cell Sci. 1998;20(1):65–70. doi:10.1023/A:1009882608636

43. Junges R, Salvadori G, Shekhar S, Åmdal HA, Periselneris JN, Chen T, et al. A Quorum-Sensing System That Regulates Streptococcus pneumoniae Biofilm Formation and Surface Polysaccharide Production. mSphere. 2017;2(5):10.1128/msphere.00324-17. doi:10.1128/msphere.00324-17

44. Morrison DA, Khan R, Junges R, Åmdal HA, Petersen FC. Genome editing by natural genetic transformation in Streptococcus mutans. Journal of Microbiological Methods. 2015;119:134–41. doi:10.1016/j.mimet.2015.09.023

45. Lemme A, Gröbe L, Reck M, Tomasch J, Wagner-Döbler I. Subpopulation-Specific Transcriptome Analysis of Competence-Stimulating-Peptide-Induced Streptococcus mutans. Journal of Bacteriology. 2011;193(8):1863–77. doi:10.1128/jb.01363-10

46. Morrison DA, Lacks SA, Guild WR, Hageman JM. Isolation and characterization of three new classes of transformation-deficient mutants of Streptococcus pneumoniae that are defective in DNA transport and genetic recombination. Journal of Bacteriology. 1983;156(1):281–90. doi:10.1128/jb.156.1.281-290.1983

47. Charpentier X, Polard P, Claverys JP. Induction of competence for genetic transformation by antibiotics: convergent evolution of stress responses in distant bacterial species lacking SOS? Current Opinion in Microbiology. 2012;Antimicrobials ° Genomics 15(5):570–6. doi:10.1016/j.mib.2012.08.001

48. Lopatkin AJ, Huang S, Smith RP, Srimani JK, Sysoeva TA, Bewick S, et al. Antibiotics as a selective driver for conjugation dynamics. Nat Microbiol. 2016;1(6):1–8. doi:10.1038/nmicrobiol.2016.44

49. Prudhomme M, Attaiech L, Sanchez G, Martin B, Claverys JP. Antibiotic Stress Induces Genetic Transformability in the Human Pathogen Streptococcus pneumoniae. Science. 2006;313(5783):89–92. doi:10.1126/science.1127912

50. Johnston C, Martin B, Fichant G, Polard P, Claverys JP. Bacterial transformation: distribution, shared mechanisms and divergent control. Nat Rev Microbiol. 2014;12(3):181–96. doi:10.1038/nrmicro3199

51. Johnston CHG, Hope R, Soulet AL, Dewailly M, De Lemos D, Polard P. The RecA-directed recombination pathway of natural transformation initiates at chromosomal replication forks in the pneumococcus. Proceedings of the National Academy of Sciences. 2023;120(8):e2213867120. doi:10.1073/pnas.2213867120

52. Overballe-Petersen S, Harms K, Orlando LAA, Mayar JVM, Rasmussen S, Dahl TW, et al. Bacterial natural transformation by highly fragmented and damaged DNA. Proceedings of the National Academy of Sciences. 2013;110(49):19860–5. doi:10.1073/pnas.1315278110

53. Kjos M, Miller E, Slager J, Lake FB, Gericke O, Roberts IS, et al. Expression of Streptococcus pneumoniae Bacteriocins Is Induced by Antibiotics via Regulatory Interplay with the Competence System. PLOS Pathogens. 2016;12(2):e1005422. doi:10.1371/journal.ppat.1005422

54. Proutière A, du Merle L, Périchon B, Varet H, Gominet M, Trieu-Cuot P, et al. Characterization of a Four-Component Regulatory System Controlling Bacteriocin Production in Streptococcus gallolyticus. mBio. 2021;12(1):10.1128/mbio.03187-20. doi:10.1128/mbio.03187-20

55. Sung CK, Morrison DA. Two Distinct Functions of ComW in Stabilization and Activation of the Alternative Sigma Factor ComX in Streptococcus pneumoniae. Journal of Bacteriology. 2005;187(9):3052–61. doi:10.1128/jb.187.9.3052-3061.2005

56. Inniss NL, Prehna G, Morrison DA. The pneumococcal σX activator, ComW, is a DNA-binding protein critical for natural transformation. J Biol Chem. 2019;294(29):11101–18. doi:10.1074/jbc.RA119.007571 PubMed PMID: 31160340; PubMed Central PMCID: PMC6643036.

57. Tovpeko Y, Bai J, Morrison DA. Competence for Genetic Transformation in Streptococcus pneumoniae: Mutations in σA Bypass the ComW Requirement for Late Gene Expression. Journal of Bacteriology. 2016;198(17):2370–8. doi:10.1128/jb.00354-16

58. Jack AA, Daniels DE, Jepson MA, Vickerman MM, Lamont RJ, Jenkinson HF, et al. Streptococcus gordonii comCDE (competence) operon modulates biofilm formation with Candida albicans. Microbiology. 2015;161(2):411–21. doi:10.1099/mic.0.000010

59. Steinmoen H, Teigen A, Håvarstein LS. Competence-Induced Cells of Streptococcuspneumoniae Lyse Competence-Deficient Cells of the SameStrain duringCocultivation. Journal of Bacteriology. 2003;185(24):7176–83. doi:10.1128/jb.185.24.7176-7183.2003

60. Cullin N, Redanz S, Lampi KJ, Merritt J, Kreth J. Murein Hydrolase LytF of Streptococcus sanguinis and the Ecological Consequences of Competence Development. Applied and Environmental Microbiology. 2017;83(24):e01709–17. doi:10.1128/AEM.01709-17

61. Moe R, Piechowiak KW, Håvarstein LS, Kjos M, Straume D. LytF contributes to pilus extrusion during natural competence in Streptococcus sanguinis SK36 [Internet]. bioRxiv; 2025 [cited 2026]. p. 2025.09.04.674207. Available from: https://www.biorxiv.org/content/10.1101/2025.09.04.674207v1 doi:10.1101/2025.09.04.674207

62. Vogel V, Fuchs M, Jachmann M, Bitzer A, Mauerer S, Münch J, et al. The Role of SilX in Bacteriocin Production of Streptococcus anginosus. Front Microbiol. 2022;13. doi:10.3389/fmicb.2022.904318

63. Nagasawa R, Ito T, Yamamoto C, Unoki M, Obana N, Nomura N, et al. Membrane vesicle production via cell-to-cell communication-induced autolysis in Streptococcus mutans. Microbiology Spectrum. 2025;13(7):e00334–25. doi:10.1128/spectrum.00334-25

64. Charpentier X, Kay E, Schneider D, Shuman HA. Antibiotics and UV Radiation Induce Competence for Natural Transformation in Legionella pneumophila. Journal of Bacteriology. 2011;193(5):1114–21. doi:10.1128/jb.01146-10

65. Sturød K, Salvadori G, Junges R, Petersen F c. Antibiotics alter the window of competence for natural transformation in streptococci. Molecular Oral Microbiology. 2018;33(5):378–87. doi:10.1111/omi.12240

66. Slager J, Kjos M, Attaiech L, Veening JW. Antibiotic-Induced Replication Stress Triggers Bacterial Competence by Increasing Gene Dosage near the Origin. Cell. 2014;157(2):395–406. doi:10.1016/j.cell.2014.01.068

67. Baddour LM, Wilson WR, Bayer AS, Fowler VG, Tleyjeh IM, Rybak MJ, et al. Infective Endocarditis in Adults: Diagnosis, Antimicrobial Therapy, and Management of Complications. Circulation. 2015;132(15):1435–86. doi:10.1161/CIR.0000000000000296

68. Lim SY, Miller JL. Ampicillin Dose for Early and Late-Onset Group B Streptococcal Disease in Neonates. Am J Perinatol. 2022;39(7):717–25. doi:10.1055/s-0040-1718880

69. Gould FK, Denning DW, Elliott TSJ, Foweraker J, Perry JD, Prendergast BD, et al. Guidelines for the diagnosis and antibiotic treatment of endocarditis in adults: a report of the Working Party of the British Society for Antimicrobial Chemotherapy. J Antimicrob Chemother. 2012;67(2):269–89. doi:10.1093/jac/dkr450

70. Dodson DS, Heizer HR, Gaensbauer JT. Sequential Intravenous-Oral Therapy for Pediatric Streptococcus anginosus Intracranial Infections. Open Forum Infect Dis. 2022;9(1):ofab628. doi:10.1093/ofid/ofab628

