## Supplementary material for "Genetic and functional characterization of the natural transformation system in *Streptococcus constellatus*"

**Table S1.** Bacterial strains, peptides, and antibiotics used in the study.

| Strain | Description | Source |
|---|---|---|
| CCUG 24889$^T$ | *Streptococcus constellatus* subsp. *constellatus* CCUG 24889 | Culture Collection University of Gothenburg |
| SC002 | CCUG 24889$^T$, but pRJ11 | This study |
| SC003 | CCUG 24889$^T$, but *psigX-fluc-aad9* (spc$^R$) | This study |
| SC005 | SC003, but Δ*comC* ::*ermB* (erm$^R$) | This study |
| SC011 | CCUG 24889$^T$, but *Δ(SCSC_RS06260 – SCSC_RS06255)* ::*aphA-3* (kan$^R$) | This study |
| SC013 | CCUG 24889$^T$, but *ΔcomC::ermB* (erm$^R$) | This study |
| SC015 | CCUG 24889$^T$, but *ΔsilED* ::*ermB* (erm$^R$) | This study |
| SC016 | SC003, but *ΔsilED* ::*ermB* (erm$^R$) | This study |
| SC017 | SC003, but *Δ(SCSC_RS06260 – SCSC_RS06255)* ::*aphA-3* (kan$^R$) | This study |
| **Peptide** | **Sequence** | **Source** |
| CSP | DSRIRMGFDFSKLFGK | GenScript, Piscataway, USA |
| **Antibiotics** | **Standard concentration** | **Manufacturer** |
| Spectinomycin | 500 µg/mL (selective plates) | Sigma-Aldrich |
| Kanamycin | 500 µg/mL (selective plates) / 50 µg/mL (reporter assay) | Sigma-Aldrich |
| Erythromycin | 10 µg/mL (selective plates) / 0.3 µg/mL (reporter assay) | Sigma-Aldrich |
| Ampicillin | 20 µg/mL (reporter assay) | Sigma-Aldrich |
| Tetracycline | 5 µg/mL (selective plates) | Sigma-Aldrich |
| Vancomycin | 4 µg/mL (reporter assay) | Sigma-Aldrich |
| Ciprofloxacin | 4 µg/mL (reporter assay) | Sigma-Aldrich |
| Chlorhexidin | 4 µg/mL (reporter assay) | Sigma-Aldrich |
| Rifampicin | 0.5 µg/mL (reporter assay) | Sigma-Aldrich |
| Chloramphenicol | 20 µg/mL (reporter assay) | Sigma-Aldrich |
| Novobiocin | 10 µg/mL (reporter assay) | Sigma-Aldrich |
| Streptomycin | 10 µg/mL (reporter assay) | Sigma-Aldrich |

**Table S2.** Donor DNA for transformation assays and oligonucleotides used in the study.

| Donor DNA | Description | Size | Primers |
|---|---|---|---|
| aSC011.3 | CCUG 24889$^{T}$ *Δ(SCSC_RS06260 – SCSC_RS06255)* ::*aphA-3* (kan$^{R}$) | 3409 bp | RJ76-77 |
| aSC011.7 | CCUG 24889$^{T}$ *Δ(SCSC_RS06260 – SCSC_RS06255)* ::*aphA-3* (kan$^{R}$) | 7097 bp | RJ108-RJ109 |
| pRJ11 | pDL278, but *aad9* (spc$^{R}$)::*aphA-3* (kan$^{R}$) | 7646 bp | |
| **Primers** | **Description** | | **5'-oligonucleotide-3'** |
| RJ23 | SC003 | F1 of SC003 | AGGAAATGATGAGCGCGAAC |
| RJ24 | | F1 of SC003 | atttacctcctcgaggatccCTTGAAGTCCATTTCATAACCC |
| RJ25 | | F2 of SC003 | gttatgaaatggacttcaagGGATCCTCGAGGAGGTAAATG |
| RJ26 | | F2 of SC003 | tactagtacagctagtggttCGCGCTTACCAATTAGAATG |
| RJ27 | | F3 of SC003 | cattctaattggtaagcgcgAACCACTAGCTGTACTAGTAG |
| RJ28 | | F3 of SC003 | TTTACGATCCGAAAACCTTC |
| RJ29 | | Nested of SC003 | GATGTGAAGGGTCGTGAAGC |
| RJ30 | | Nested of SC003 | AGATTCCCTACTGCTGCCTC |
| RJ60 | SC005 | F1 of SC005 | CGTACAATGATGGGCTGAATAATC |
| RJ61 | | F1 of SC005 | tttgggcccgGCTAAAAATTTTTTTCATAAATTCTCCTTAATG |
| RJ62 | | F2 of SC005 | aatttttagcCGGGCCCAAAATTTGTTTGATTTG |
| RJ63 | | F2 of SC005 | aatcaaatccGCGACTCATAGAATTATTTCCTCC |
| RJ64 | | F3 of SC005 | tatgagtcgcGGATTTGATTTTTCTAAACTTTTTGG |
| RJ65 | | F3 of SC005 | CTTTTTGCTCAGTTTGGGTTTTATTATATC |
| RJ66 | | Nested of SC005 | AGGATTTGTTTGGATGTCATTG |
| RJ67 | | Nested of SC005 | GCACCCTTTATGAATGCTTAGT |
| RJ68 | | Detect comC | GCAAAAAGGAAATAACTAAAGTGGA |
| RJ69 | | Detect comC | TCAAATCCCATTCTTATTCGAC |
| RJ70 | SC011 | F1 of SC011 | TTCAAGTGCGGGGAAATTG |
| RJ71 | | F1 of SC011 | aaacggcgcgATGATTATCAGCAAGTTTTTCTTG |
| RJ72 | | F2 of SC011 | tgataatcatCGCGCCGTTTGATTTTTAATG |
| RJ73 | | F2 of SC011 | ctttttctccCCGGCCATCGATACAAATTC |
| RJ74 | | F3 of SC011 | cgatggccggGGAGAAAAAGAGGAGCTAAAAATAAC |
| RJ75 | | F3 of SC011 | AACAGAGCACAAGTTGAAC |
| RJ76 | | Nested of SC011.3kb | GAAATTGGGAAGAtAGcATTTGG |
| RJ77 | | Nested of SC011.3kb | AATGATAATCGTTTCATTGTCTGC |
| RJ108 | SC011 | Nested of SC011.7kb | CCTTTTGTCGGTACATAAGAAGA |
| RJ109 | | Nested of SC011.7kb | CTCAGAAATTAGCAACCATTATGC |
| RJ110 | SC016 | F1 of SC016 | CTGCCAGAGAATAATATGC |
| RJ111 | | F1 of SC016 | tttgggcccgGCAATCTCTTGCATCTATTTG |
| RJ112 | | F2 of SC016 | aagagattgcCGGGCCCAAAATTTGTTTG |
| RJ113 | | F2 of SC016 | tgacaatagtGCGACTCATAGAATTATTTCCTC |
| RJ114 | | F3 of SC016 | tatgagtcgcACTATTGTCATTGCGAAAAAATC |
| RJ115 | | F3 of SC016 | GATATCGCAGCTTTTCTTAC |
| RJ116 | | F4 of SC016 | TTTATCCACCCCAGCGAAC |
| RJ117 | | F4 of SC016 | AACTTTATAGTAAGGTTGGTATCTGGA |
| RJ128 | *coiA* | F | AAGGAAGAGTCATGCGACCG |
| RJ129 | | R | GCCCAAGTAAACAAGGCAGC |
| RJ130 | *ciaH* | F | ACTCTTTGTCAGCTGTTCCACT |
| RJ131 | | R | TCAAGTTATGCGCTCAAGCC |
| RJ136 | *cclA* | F | GCAACGCAGCCGATTGATAA |
| RJ137 | | R | TGGTCGTTGATCGCTTTCCT |
| RJ140 | *comEA* | F | ATCCAAAAAGCGGGTGGACT |
| RJ141 | | R | ATTCCCTTTTTGTGCCGCAG |
| RJ142 | *dprA* | F | TCAATTCCGCGTGCTAAACC |
| RJ143 | | R | TAGCCGAGTAGGCAGTCAATC |
| RJ144 | SCSC_RS09450 / hyp. prot. | F | TCTGACACAATCTTCATCTTTTGC |
| RJ145 | | R | ACAGAAGTCATCACCGTTTTCG |
| RJ146 | *comGA* | F | GGCGTTCATCTCCAATTCGC |
| RJ147 | | R | GGAAGAACGAGCACAGGACA |
| RJ148 | *ssbB* | F | CGGCTAATTTGCCCCAAACA |
| RJ149 | | R | GCAACGCTTGCTGTTAATCG |
| RJ150 | *cinA* | F | CCAGTCGACTCGCGAAGAAA |
| RJ151 | | R | GTGGCTTGGGTCCAACAGAA |
| RJ156 | *comFA* | F | GCAGTTGCCAAGCGAAAAGA |
| RJ157 | | R | TCGCCATGCAGGAGTGAAAT |
| RJ160 | *dut* | F | GGCGCAATAACTGTACGCTC |
| RJ161 | | R | TACTGCCAAAGCGTGAGACG |
| RJ164 | *pheT* | F | TAGCATATGAGGTGGCAGCG |
| RJ165 | | R | ACGCACTGCATAAAAAGGCG |
| RJ168 | *radC* | F | TTGCCAAGAGAACGTTTGGT |
| RJ169 | | R | GCCAGTGCGGATAAGAATGG |
| RJ190 | *gyrA* | F | GCACGAGCTTCTGCCTTTTC |
| RJ191 | | R | TCCTAAAATTCTCTCCTTGCGTGA |
| RJ204 | *lytF* | F | GCAGCATAATTGGGTTGAAAGG |
| RJ205 | | R | CTGGTCTGAAGTCAACGGGC |
| RJ206 | *comD* | F | TCCCTCGCCCTTTGTTGAAA |
| RJ207 | | R | TGCCATAGAAGGAGCAGCAG |
| RJ208 | *sigX* | F | TTGGGACCAAGAAGGGATGC |
| RJ209 | | R | GCTTTCTTGTTTGCGCAGGA |
| RJ210 | *murB* | F | CAAAACCGCGAATACCACCG |
| RJ211 | | R | GGTGGAGCTGCCGATTATCT |
| RJ214 | *spx* | F | CATTACGAACCGAGCGAGGTA |
| RJ215 | | R | TGGGGTTGCAAGTATTGTGTCT |
| RJ216 | SCSC_RS05470 (ORF-3) | F | GTGAAAGAGGCGAGCGTTTG |
| RJ217 | | R | AGAGTTAGCGAGGCGCTTTC |
| RJ218 | SCSC_RS05830 (ORF-1) | F | TGAGTTCGCCAATGGAAGTGT |
| RJ219 | | R | ATTGGTCGGTGCGTGGTTTA |
| RJ220 | SCSC_RS05940 (ORF-3) | F | ATACCCACATGGCACACCAG |
| RJ221 | | R | CCAATGGAGGTATCGGAGGC |
| RJ222 | *murC* (ORF-2) | F | GGAACCCGTCGTAGAACGAA |
| RJ223 | | R | ACAAGGAGTTGCGTCGGATT |
| RJ230 | *ciaR* | F | AATAATCGTCGGCTCCCAGC |
| RJ231 | | R | TACGAAGCCGAAAGTGGTGT |

UPPERCASE: Annealing region, lowercase: overlapping region.

**Table S3.** Genomes utilized in the study.

| Species | Sub-species | Strain | Genome size [bp] | G + C [%] | CDS | rRNA | Pseudogenes | Level | Accession |
|---|---|---|---|---|---|---|---|---|---|
| *S. constellatus* | *constellatus* | SK53 | 1 840 061 | 38 | 1 715 | 38 | 80 | Contig | GCF_000257785.1 |
| *S. constellatus* | *pharyngis* | C232 | 1 935 414 | 38 | 1 790 | 74 | 127 | Complete | GCF_000463395.1 |
| *S. constellatus* | *pharyngis* | C1050 | 1 991 156 | 38 | 1 839 | 74 | 127 | Complete | GCF_000463425.1 |
| *S. constellatus* | *pharyngis* | C818 | 1 935 662 | 38 | 1 787 | 74 | 129 | Complete | GCF_000463445.1 |
| *S. constellatus* | *pharyngis* | CCUG 46377 | 1 950 566 | 38 | 1 807 | 53 | 159 | Contig | GCF_000474135.1 |
| *S. constellatus* | | KCOM 1650 | 1 965 746 | 38 | 1 837 | 49 | 80 | Contig | GCF_000814045.1 |
| *S. constellatus* | | 317_SINT | 1 853 071 | 38 | 1 712 | 54 | 86 | Contig | GCF_001072275.1 |
| *S. constellatus* | | 783_SANG | 1 913 462 | 38 | 1 809 | 37 | 85 | Scaffold | GCF_001074375.1 |
| *S. constellatus* | | 925_SCON | 2 043 273 | 38 | 1 965 | 35 | 101 | Contig | GCF_001075725.1 |
| *S. constellatus* | | KCOM 1039 | 1 885 802 | 38 | 1 728 | 71 | 83 | Scaffold | GCF_003570855.1 |
| *S. constellatus* | *constellatus* | ATCC 27823 | 1 867 902 | 38 | 1 764 | 49 | 85 | Scaffold | GCF_008633005.1 |
| *S. constellatus* | *constellatus* | 13-11-14 | 1 800 973 | 38 | 1 698 | 57 | 79 | Scaffold | GCF_008633025.1 |
| *S. constellatus* | *pharyngis* | 15-01-28 | 1 914 344 | 38 | 1 791 | 62 | 125 | Scaffold | GCF_008633405.1 |
| *S. constellatus* | | D6t1_180914_C10 | 1 863 546 | 38 | 1 756 | 43 | 90 | Scaffold | GCF_015555235.1 |
| *S. constellatus* | | 1001254J_160919_C10 | 1 755 446 | 38 | 1 627 | 48 | 82 | Scaffold | GCF_015559305.1 |
| *S. constellatus* | | FDAARGOS_1015 | 2 038 583 | 38 | 1 875 | 73 | 113 | Complete | GCF_016127875.1 |
| *S. constellatus* | | FDAARGOS_1156 | 1 903 262 | 38 | 1 741 | 74 | 83 | Complete | GCF_016725005.1 |
| *S. constellatus* | | FDAARGOS_1208 | 1 978 680 | 38 | 1 827 | 74 | 124 | Complete | GCF_016889885.1 |
| *S. constellatus* | | 1540534 | 1 869 421 | 38 | 1 757 | 49 | 106 | Contig | GCF_019336935.1 |
| *S. constellatus* | | S60 | 1 892 624 | 38 | 1 772 | 47 | 90 | Scaffold | GCF_023109175.1 |
| *S. constellatus* | | S54 | 1 927 926 | 38 | 1 827 | 49 | 85 | Scaffold | GCF_023109215.1 |
| *S. constellatus* | | S55 | 1 993 590 | 38 | 1 867 | 48 | 90 | Scaffold | GCF_023109295.1 |
| *S. constellatus* | | S34 | 1 818 626 | 38 | 1 734 | 39 | 88 | Scaffold | GCF_023109615.1 |
| ***S. constellatus*** | ***constellatus*** | **CCUG 24889** | **1 901 581** | **38** | **1 738** | **74** | **84** | **Complete** | **GCF_023167545.1** |
| *S. constellatus* | | TCV107 | 1 980 997 | 38 | 1 828 | 74 | 101 | Complete | GCF_024399395.1 |
| *S. constellatus* | | AM109-96 | 1 821 433 | 38 | 1 700 | 50 | 69 | Scaffold | GCF_027723325.1 |
| *S. constellatus* | | UMB8371 | 1 830 535 | 38 | 1 704 | 30 | 79 | Contig | GCF_030215825.1 |
| *S. constellatus* | | SMC7155 | 1 826 913 | 38 | 1 694 | 48 | 92 | Contig | GCF_030676655.1 |
| *S. constellatus* | | 1033st1_A6_1033SCRN_220408 | 1 934 643 | 38 | 1 822 | 40 | 81 | Scaffold | GCF_039059945.1 |
| *S. constellatus* | | 1033st1_F12_1033SCRN_220408 | 1 925 965 | 38 | 1 816 | 48 | 78 | Scaffold | GCF_039059995.1 |
| *S. constellatus* | | 20925_1_21 | 1 949 580 | 38 | 1 812 | 39 | 87 | Contig | GCF_041432565.1 |
| *S. constellatus* | | NCTC11325 | 1 906 855 | 38 | 1 743 | 74 | 86 | Contig | GCF_900459125.1 |
| *S. constellatus* | | SS_Bg39 | 2 040 680 | 39 | 1 937 | 57 | 92 | Contig | GCF_902167705.1 |

**Table S4.** List and source of gene queries utilized in the study.

| Gene Name | Locus Tag | Organism | Accession | Gene Start | Gene Stop |
|---|---|---|---|---|---|
| *comA* | AT689_RS02160 | *S. pneumoniae* NCTC7465 | NZ_LN831051.1 | 418894 | 421047 |
| *comB* | AT689_RS02155 | *S. pneumoniae* NCTC7465 | NZ_LN831051.1 | 417532 | 418881 |
| *comC* $_{SP}$ | AT689_RS02690 | *S. pneumoniae* NCTC7465 | NZ_LN831051.1 | 500327 | 500452 |
| *comD* | AT689_RS02695 | *S. pneumoniae* NCTC7465 | NZ_LN831051.1 | 500473 | 501798 |
| *comE* | AT689_RS02700 | *S. pneumoniae* NCTC7465 | NZ_LN831051.1 | 501795 | 502547 |
| *comW* $_{SP}$ | AT689_RS02540 | *S. pneumoniae* NCTC7465 | NZ_LN831051.1 | 474769 | 475005 |
| *comW* $_{SA}$ | FGK97_RS09015 | *S. anginosus* NCTC11064 | NZ_LR594037.1 | 1782424 | 1782654 |
| *comX1* | AT689_RS02600 | *S. pneumoniae* NCTC7465 | NZ_LN831051.1 | 482441 | 482920 |
| *comX2* | AT689_RS03915 | *S. pneumoniae* NCTC7465 | NZ_LN831051.1 | 741124 | 741603 |
| *comFA* | AT689_RS02850 | *S. pneumoniae* NCTC7465 | NZ_LN831051.1 | 528548 | 529846 |
| *comFC* | AT689_RS02855 | *S. pneumoniae* NCTC7465 | NZ_LN831051.1 | 529843 | 530505 |
| *comEA* | AT689_RS09050 | *S. pneumoniae* NCTC7465 | NZ_LN831051.1 | 1710010 | 1710660 |
| *comEC* | AT689_RS09045 | *S. pneumoniae* NCTC7465 | NZ_LN831051.1 | 1707786 | 1710026 |
| *comGA* | AT689_RS03685 | *S. pneumoniae* NCTC7465 | NZ_LN831051.1 | 701239 | 702180 |
| *comGB* | AT689_RS03690 | *S. pneumoniae* NCTC7465 | NZ_LN831051.1 | 702128 | 703144 |
| *comGC* | AT689_RS03695 | *S. pneumoniae* NCTC7465 | NZ_LN831051.1 | 703146 | 703472 |
| *comGD* | AT689_RS03700 | *S. pneumoniae* NCTC7465 | NZ_LN831051.1 | 703465 | 703869 |
| *comGE* | AT689_RS03705 | *S. pneumoniae* NCTC7465 | NZ_LN831051.1 | 703832 | 704134 |
| *comGF* | AT689_RS03710 | *S. pneumoniae* NCTC7465 | NZ_LN831051.1 | 704097 | 704558 |
| *comGG* | AT689_RS03715 | *S. pneumoniae* NCTC7465 | NZ_LN831051.1 | 704536 | 704949 |
| *cinA* | AT689_RS04290 | *S. pneumoniae* NCTC7465 | NZ_LN831051.1 | 804272 | 805528 |
| *coiA* | AT689_RS08930 | *S. pneumoniae* NCTC7465 | NZ_LN831051.1 | 1687997 | 1688950 |
| *cclA* | AT689_RS05025 | *S. pneumoniae* NCTC7465 | NZ_LN831051.1 | 926954 | 927613 |
| *endA* | AT689_RS04210 | *S. pneumoniae* NCTC7465 | NZ_LN831051.1 | 784635 | 785459 |
| *radA* | AT689_RS02250 | *S. pneumoniae* NCTC7465 | NZ_LN831051.1 | 434582 | 435943 |
| *recA* | AT689_RS04295 | *S. pneumoniae* NCTC7465 | NZ_LN831051.1 | 805583 | 806749 |
| *dprA* | AT689_RS07605 | *S. pneumoniae* NCTC7465 | NZ_LN831051.1 | 1415325 | 1416173 |
| *ssbB* | AT689_RS04440 | *S. pneumoniae* NCTC7465 | NZ_LN831051.1 | 826850 | 827245 |
| *qsrA* | AT689_RS05575 | *S. pneumoniae* NCTC7465 | NZ_LN831051.1 | 1036194 | 1037087 |
| *cbpA* | SPD_RS10670 | *S. pneumoniae* D39 | NC_008533.2 | 1995045 | 1997150 |
| *cbpB* | SPD_RS08840 | *S. pneumoniae* D39 | NC_008533.2 | 1670508 | 1670969 |
| *cbpC* | SPD_RS01875 | *S. pneumoniae* D39 | NC_008533.2 | 349486 | 350502 |
| *cbpD* | SPD_RS10730 | *S. pneumoniae* D39 | NC_008533.2 | 2006513 | 2007859 |
| *cbpE* | SPD_RS04410 | *S. pneumoniae* D39 | NC_008533.2 | 838512 | 840395 |
| *cbpF* | SPD_RS01940 | *S. pneumoniae* D39 | NC_008533.2 | 358923 | 359807 |
| *cbpG* | SPD_RS01935 | *S. pneumoniae* D39 | NC_008533.2 | 358079 | 358904 |
| *lytA* | AT689_RS04305 | *S. pneumoniae* NCTC7465 | NZ_LN831051.1 | 808802 | 809758 |
| *lytC* | AT689_RS06215 | *S. pneumoniae* NCTC7465 | NZ_LN831051.1 | 1159820 | 1161325 |
| *lytF* $_{SG}$ | SGO_RS10250 | *S. gordonii* CH1 | NC_009785.1 | 2155996 | 2157645 |
| *lytF* $_{SA}$ | HMPREF9966_RS05520 | *S. anginosus* SK52 | NZ_AFIM01000066.1 | 27278 | 29212 |
| *comM* | AT689_RS04270 | *S. pneumoniae* NCTC7465 | NZ_LN831051.1 | 801506 | 802126 |
| *def2* | AT689_RS06330 | *S. pneumoniae* NCTC7465 | NZ_LN831051.1 | 1183011 | 1183421 |
| *dut* | AT689_RS02260 | *S. pneumoniae* NCTC7465 | NZ_LN831051.1 | 436474 | 436917 |
| *blpA* | AT689_RS11025 | *S. pneumoniae* NCTC7465 | NZ_LN831051.1 | 2082276 | 2084429 |
| *blpB* | AT689_RS11030 | *S. pneumoniae* NCTC7465 | NZ_LN831051.1 | 2084440 | 2085801 |
| *blpC* $_{SP}$ | AT689_RS11035 | *S. pneumoniae* NCTC7465 | NZ_LN831051.1 | 2085858 | 2086013 |
| *blpC* $_{SA}$ | FGK97_RS02595 | *S. anginosus* NCTC11064 | NZ_LR594037.1 | 510497 | 510646 |
| *blpH* $_{SP}$ | AT689_RS11040 | *S. pneumoniae* NCTC7465 | NZ_LN831051.1 | 2086057 | 2087397 |
| *blpH* $_{SA}$ | FGK97_RS02600 | *S. anginosus* NCTC11064 | NZ_LR594037.1 | 510691 | 511998 |
| *blpR* | AT689_RS11045 | *S. pneumoniae* NCTC7465 | NZ_LN831051.1 | 2087411 | 2088148 |
| *blp3.1* | | *S. anginosus* BSU 1211 | MZ766502.1 | 286 | 516 |
| *blp3.2* | | *S. anginosus* BSU 1211 | MZ766502.1 | 746 | 1066 |
| *blp3.3* | | *S. anginosus* BSU 1211 | MZ766502.1 | 1218 | 1508 |
| *blp3.4* | | *S. anginosus* BSU 1211 | MZ766502.1 | 1560 | 1814 |
| *blp3.5* | | *S. anginosus* BSU 1211 | MZ766502.1 | 1801 | 2187 |
| *blp3.6* | | *S. anginosus* BSU 1211 | MZ766502.1 | 2209 | 2433 |
| *silX* | | *S. anginosus* BSU 1211 | MZ766502.1 | 2665 | 3378 |
| *silA* | | *S. anginosus* BSU 1211 | MZ766502.1 | 3541 | 4293 |
| *silB* | | *S. anginosus* BSU 1211 | MZ766502.1 | 4298 | 5605 |
| *silCR* | | *S. anginosus* BSU 1211 | MZ766502.1 | 5650 | 5799 |
| *silD* | | *S. anginosus* BSU 1211 | MZ766502.1 | 5892 | 7253 |
| *silE* | | *S. anginosus* BSU 1211 | MZ766502.1 | 7264 | 9414 |

Footnotes in gene name indicate the same gene target using gene sequences from different species

$_{SP}$: *Streptococcus pneumoniae* . $_{SA}$: *Streptococcus anginosus* . $_{SG}$: *Streptococcus gordonii* .

**Table S5.** Comparative analysis of exporter similarity.

| Species | Subsp. | Strain | *comA* SP | *blpA* SP | *silE* SA | *comB* SP | *blpB* SP | *silD* SA |
|---|---|---|---|---|---|---|---|---|
| *Streptococcus constellatus* | | 1001254J_160919_C10 | 70.59% | 82.72% | 87.56% | 42.28% | 59.51% | 80.92% |
| *Streptococcus constellatus* | | 1033st1_A6_1033SCRN_220408 | 70.78% | 83.75% | 89.90% | 42.88% | 60.07% | 81.84% |
| *Streptococcus constellatus* | | 1033st1_F12_1033SCRN_220408 | 70.78% | 83.75% | 89.90% | 42.88% | 60.07% | 81.84% |
| *Streptococcus constellatus* | | 11-6117 | 70.95% | 83.92% | 90.34% | 42.92% | 60.29% | 82.45% |
| *Streptococcus constellatus* | | 20925_1_21 | 70.96% | 83.93% | 90.16% | 42.56% | 59.92% | 82.44% |
| *Streptococcus constellatus* | | 317_SINT | 70.39% | 83.25% | 89.65% | 42.59% | 60.18% | 81.93% |
| *Streptococcus constellatus* | | 783_SANG | 70.96% | 83.93% | 90.16% | 42.56% | 59.92% | 82.44% |
| *Streptococcus constellatus* | | 925_SCON | 70.74% | 83.65% | 90.07% | 42.47% | 59.86% | 81.78% |
| *Streptococcus constellatus* | | AM109-96 | 70.63% | 83.89% | 89.92% | 42.70% | 60.25% | 82.19% |
| *Streptococcus constellatus* | | D6t1_180914_C10 | 70.80% | 82.93% | 87.77% | 42.32% | 59.51% | 80.70% |
| *Streptococcus constellatus* | | FDAARGOS_1015 | 70.34% | 83.37% | 89.43% | 42.51% | 60.24% | 82.44% |
| *Streptococcus constellatus* | | FDAARGOS_1156 | 70.83% | 83.39% | 88.06% | 42.62% | 60.48% | 82.25% |
| *Streptococcus constellatus* | | FDAARGOS_1208 | 70.37% | 83.41% | 89.47% | 42.51% | 60.24% | 82.44% |
| *Streptococcus constellatus* | | KCOM 1039 | - | - | - | 42.45% | 60.12% | 82.18% |
| *Streptococcus constellatus* | | KCOM 1650 | 70.95% | 83.92% | 90.34% | 42.53% | 60.53% | 82.08% |
| *Streptococcus constellatus* | | NCTC11325 | 70.83% | 83.39% | 88.06% | 42.57% | 60.43% | 82.19% |
| *Streptococcus constellatus* | | S34 | 70.72% | 83.55% | 89.89% | 41.87% | 59.23% | 80.81% |
| *Streptococcus constellatus* | | S54 | 70.29% | 83.34% | 89.95% | 42.83% | 59.99% | 82.09% |
| *Streptococcus constellatus* | | S55 | 70.36% | 83.63% | 89.74% | 42.57% | 60.34% | 81.88% |
| *Streptococcus constellatus* | | S60 | 70.91% | 84.13% | 90.36% | 42.61% | 59.80% | 82.19% |
| *Streptococcus constellatus* | | SMC7155 | 70.83% | 83.39% | 88.06% | 42.62% | 60.48% | 82.25% |
| *Streptococcus constellatus* | | SS_Bg39 | 71.26% | 84.02% | 95.68% | 42.23% | 60.74% | 92.72% |
| *Streptococcus constellatus* | *constellatus* | 13-11-14 | 70.79% | 83.85% | 90.16% | 42.62% | 60.22% | 82.21% |
| *Streptococcus constellatus* | *constellatus* | ATCC 27823 | 70.67% | 83.23% | 91.47% | 42.83% | 60.24% | 82.57% |
| ***Streptococcus constellatus*** | ***constellatus*** | **CCUG 24889** | **70.07%** | **83.21%** | **89.76%** | **42.63%** | **60.10%** | **82.27%** |
| *Streptococcus constellatus* | *constellatus* | SK53 | 70.83% | 83.39% | 88.06% | 42.62% | 60.48% | 82.25% |
| *Streptococcus constellatus* | *pharyngis* | 15-01-28 | 70.83% | 83.39% | 88.06% | 42.62% | 60.48% | 82.25% |
| *Streptococcus constellatus* | *pharyngis* | C1050 | 70.83% | 83.39% | 88.06% | 42.62% | 60.48% | 82.25% |
| *Streptococcus constellatus* | *pharyngis* | C232 | 70.43% | 83.46% | 89.32% | 42.51% | 60.24% | 82.44% |
| *Streptococcus constellatus* | *pharyngis* | C818 | 70.37% | 83.41% | 89.47% | 42.51% | 60.24% | 82.44% |
| *Streptococcus constellatus* | *pharyngis* | SK1060 | 70.37% | 83.41% | 89.47% | 40.81% | 58.34% | 79.42% |
| *Streptococcus constellatus* | | TCV107 | 70.37% | 83.41% | 89.47% | 40.81% | 58.34% | 79.42% |
| *Streptococcus constellatus* | | UMB8371 | 70.37% | 83.41% | 89.47% | 42.51% | 60.24% | 82.44% |

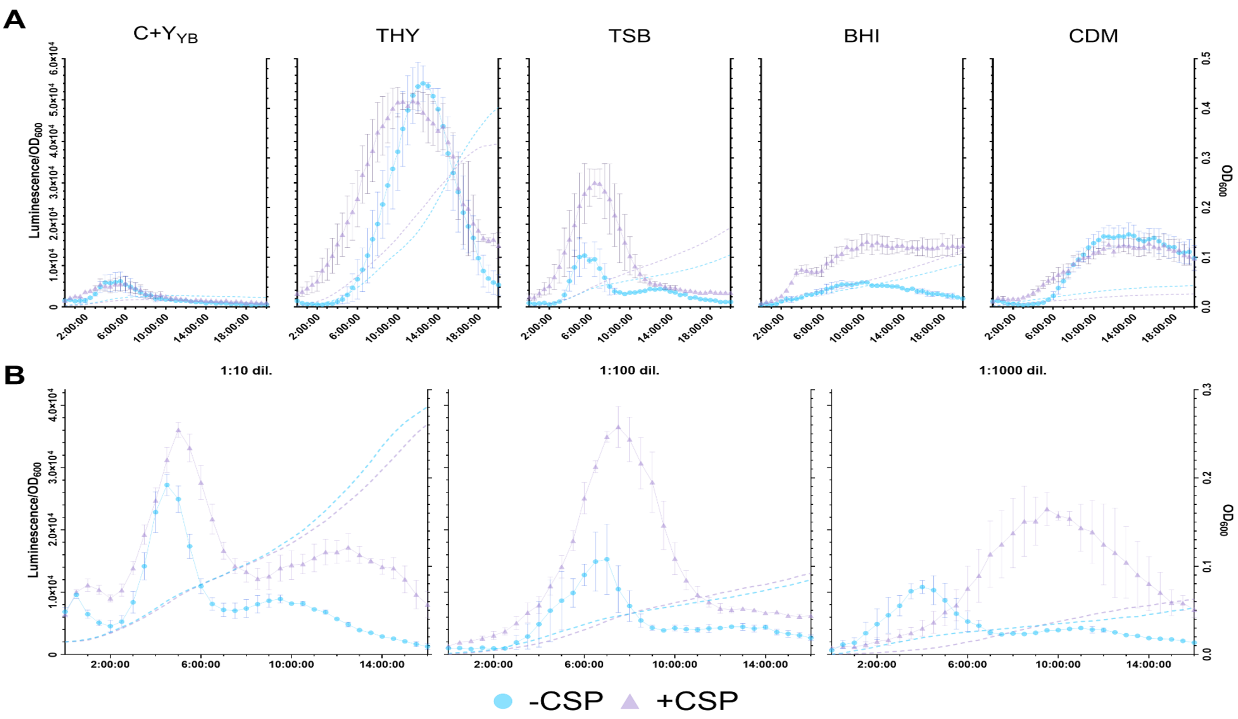


**Fig S1.** Influence of media and dilution factors on *sigX* expression. **(A)** Comparative *sigX* expression profiles across different growth media ($C+Y_{YB}$, THY, TSB, BHI, and CDM) ±CSP. **(B)** Effect of various inoculation dilutions (1:10, 1:100, and 1:1000) in TSB on the timing and intensity of the competence response. All data represent mean ± SEM of three independent biological experiments.
